# A mechanistic model captures the emergence and implications of non-genetic heterogeneity and reversible drug resistance in ER+ breast cancer cells

**DOI:** 10.1101/2021.03.14.435359

**Authors:** Sarthak Sahoo, Ashutosh Mishra, Harsimran Kaur, Kishore Hari, Srinath Muralidharan, Susmita Mandal, Mohit Kumar Jolly

## Abstract

Resistance to anti-estrogen therapy is an unsolved clinical challenge in successfully treating ER+ breast cancer patients. Acquisition of mutations can confer heritable resistance to cancer cells, enabling their clonal selection to establish a drug-resistant population. Recent studies have demonstrated that cells can tolerate drug treatment without any genetic alterations too; however, the mechanisms and dynamics of such non-genetic adaptation remain elusive. Here, we investigate coupled dynamics of epithelial-mesenchymal transition (EMT) in breast cancer cells and emergence of reversible drug resistance. Our mechanism-based model for the underlying regulatory network reveals that these two axes can drive one another, thus conferring bidirectional plasticity. This network can also enable non-genetic heterogeneity in a population of cells by allowing for six co-existing phenotypes: epithelial-sensitive, mesenchymal-resistant, hybrid E/M-sensitive, hybrid E/M-resistant, mesenchymal-sensitive and epithelial-resistant, with the first two ones being most dominant. Next, in a population dynamics framework, we exemplify the implications of phenotypic plasticity (both drug-induced and intrinsic stochastic switching) and/or non-genetic heterogeneity in promoting population survival in a mixture of sensitive and resistant cells, even in the absence of any cell-cell cooperation. Finally, we propose the potential therapeutic use of MET (mesenchymal-epithelial transition) inducers besides canonical anti-estrogen therapy to limit the emergence of reversible drug resistance. Our results offer mechanistic insights into empirical observations on EMT and drug resistance and illustrate how such dynamical insights can be exploited for better therapeutic designs.

## Introduction

Emergence of drug resistance remains the biggest hurdle in clinical management of cancer. It has been largely tacitly assumed that the acquisition of genomic mutations is a necessary and sufficient condition for drug resistance. However, recent studies across multiple cancers have suggested a set of alternative non-genetic mechanisms that can facilitate the survival of cancer cells in the presence of cytotoxic therapies (1). These non-genetic mechanisms do not entail changes in genotype (underlying DNA sequence), but in the manifestation of phenotype (2, 3) through epigenetic or transcriptional reprogramming and/or cell-state transitions (4–8). Unlike genetic changes which are ‘hard-wired’ and irreversibly passed to further generations, the non-genetic changes are reversible and stochastic in nature and thus not necessarily heritable. None of the existing therapies has been yet shown to be capable of outsmarting this adaptive ability of cancer cells to alter their phenotype without modifying their genotype. Instead, drug treatment can promote such cell-state transitions, thus potentially worsening the disease progression (9). Thus, despite major advancements in targeted therapy, mechanisms of non-genetic heterogeneity and reversible drug resistance remain largely elusive.

Tamoxifen was the first targeted therapy for breast cancer which was given to ER+ (Estrogen receptor-positive) breast cancer patients to bind to Estrogen Receptor (ER) and antagonize the proliferative ability potentiated by binding of ER to growth hormone estrogen (10). Estrogen receptor alpha (ERα) is one of the two forms of ER that lies upstream to various genomic and non-genomic signalling pathways that control cellular proliferation and survival, essentially regulating the growth of normal breast tissue and tumor (10). ERα is considered as a key prognostic marker; increased response to anti-estrogen therapies (such as tamoxifen) and better patient survival is associated with higher levels of ERα (11). However, one-third of women treated with tamoxifen for five years have recurrent disease within 15 years, with a majority of them being metastatic (10, 12). Thus, development of acquired resistance to tamoxifen limits its efficacy.

One of the molecules associated with tamoxifen insensitivity is ER-α36, a variant form of ERα (also known as ER-α66) (13, 14). As compared to ER-α66, ER-α36 lacks both transcriptional activation domains (AF-1 and AF-2) but retains the DNA-binding and ligand-binding domains. ER-α36 has been associated with activating downstream signaling pathways that promote cell proliferation, highlighting its potential role in development of drug resistance against anti-estrogen treatment (15–17). Mechanistically, ER-α36 has been shown to be transcriptionally activated by tamoxifen (18) but inhibited by ER-α66 (19).

Another process associated with tamoxifen resistance is epithelial-mesenchymal transition (EMT) (20, 21), a cell plasticity program involved with drug resistance across cancers (22, 23). Loss of ERα induced changes concomitant with EMT (24). Consistently, tamoxifen-resistant cells were seen to grow loose colonies with weak cell-cell adhesion, typical of EMT (25). On the other hand, EMT-inducing transcription factors such as SLUG, ZEB1 and SNAIL are known to drive tamoxifen resistance (26–28). This bidirectional coupling between ERα and EMT pathways is reinforced by analysis of 118 breast tumor specimens showing specific association of ER-α36 with various EMT markers such as MMP9, SNAIL1 and VIM (29). However, given that EMT is a reversible phenomenon where cancer cells can stably acquire one or more hybrid epithelial/ mesenchymal phenotypes too (30), many questions remain: a) can EMT drive acquisition of reversible resistance to tamoxifen, i.e. can the reverse of EMT – Mesenchymal-Epithelial Transition (MET) – restore tamoxifen sensitivity? b) can tamoxifen resistance drive a partial or full EMT? and c) do cells need to undergo a full EMT to gain tamoxifen resistance, or can epithelial and hybrid E/M cells also possibly show those traits?

Here, we develop a mechanism-based model based on a gene regulatory network among known between components of EMT and tamoxifen resistance (TamR) associated pathways. Dynamical simulations of this network reveals different “attractors” (expression patterns) that can emerge, thus enabling the (co)-existence of various states along EMT and TamR axes: ES (epithelial – Tam sensitive), ER (epithelial – Tam resistant), HS (hybrid – Tam sensitive), HR (hybrid – Tam resistant), MS (mesenchymal – Tam sensitive) and MR (mesenchymal – Tam resistant). Further, the emergent dynamics of the coupled processes of EMT and TamR facilitates either process to be able to drive another one, thus enabling cells to switch among multiple phenotypes along these interconnected axes and driving non-genetic heterogeneity. Finally, we develop a population dynamics model to decipher the contribution of intrinsic and tamoxifen-induced phenotypic plasticity and non-genetic heterogeneity in a cell population. Our simulations suggest that the long-term maintenance of TamR cells in a population can arise from many possible scenarios: a) non-genetic heterogeneity in sensitivity to tamoxifen in the initial cell population, b) phenotypic plasticity enabled by EMT and/or drug treatment. Thus, the emergence of TamR can be potentially curtailed through combinatorial targeting both EMT and ERα pathways.

## Results

### Crosstalk between EMT and Estrogen receptor signaling result in multiple phenotypes showing broad association between EMP and drug resistance in ER+ breast cancer

First, we identified a gene regulatory network (GRN) that captures known interactions among various players involved in EMT and in tamoxifen resistance. This network incorporates the reported interactions among estrogen receptor molecules ERα66 and ERα36, and EMT regulators SLUG, ZEB1 and miR-200 (**Fig 1A**). ERα66 can repress ERα36 expression in an estrogen-independent manner (19), and activate its own expression (31). ERα66 can also exert controls over the EMT axis by repressing SLUG (32). SLUG and ZEB1 are key EMT-inducing transcription factors that can regulate the expression of ERα66 and ERα36, thereby controlling tamoxifen resistance. SLUG and ZEB1 can repress ERα66 (28, 33), while ERα36 can enhances the expression of ZEB1 through SNAIL and/or by suppression of CDH1 (18, 34, 35). ZEB1 and miR-200 form a mutually inhibitory self-activatory feedback loop, and in conjunction with their interaction with SLUG, they determine the EMT phenotype of a cell (35).

**Figure 1:**
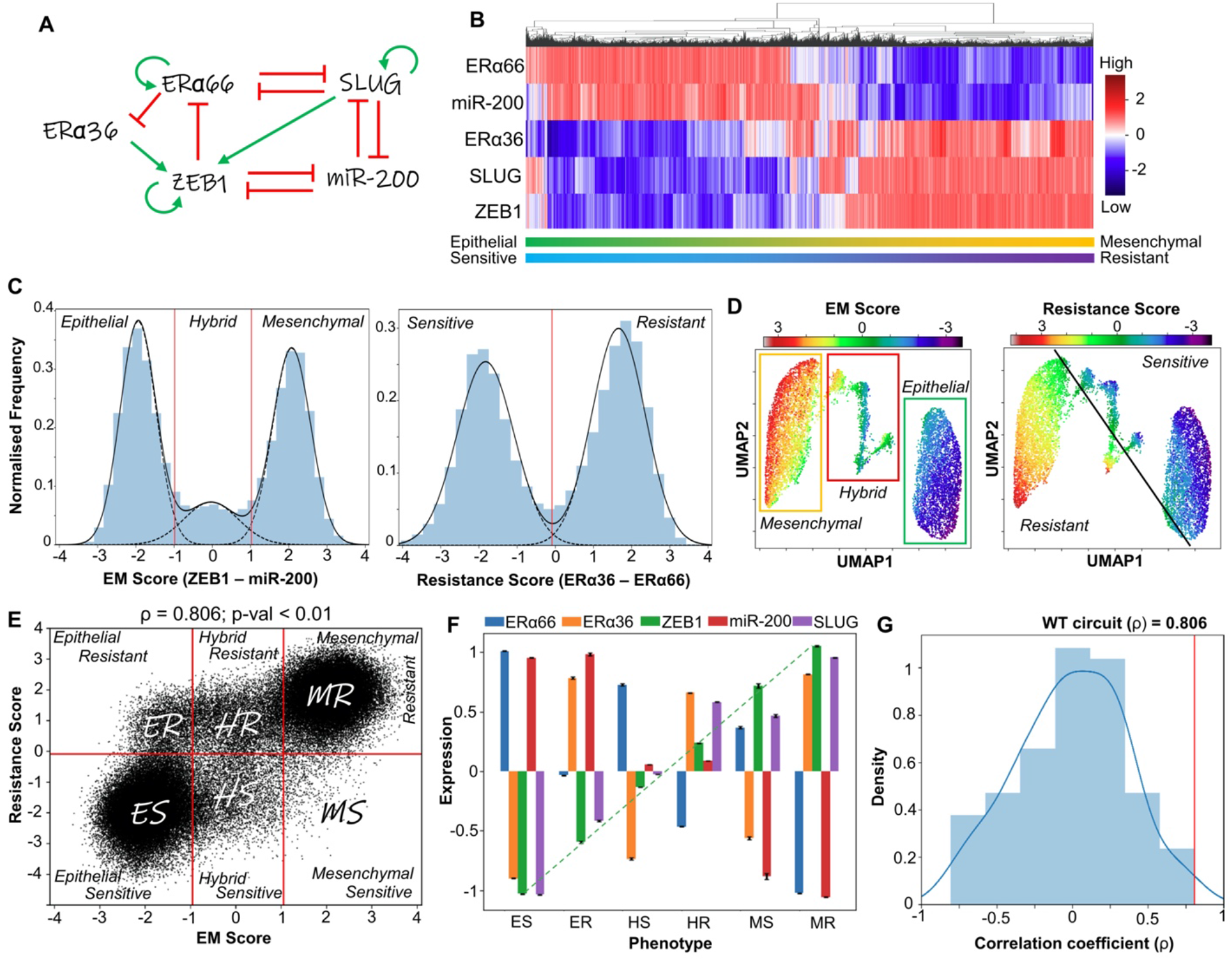
Emergent dynamics of coupled EMT-ERα signaling network. **A**. Gene regulatory network (GRN) showing crosstalk between EMT and Estrogen receptor (ERα) signaling. Green arrows represent activation links; red hammers represent inhibition. **B**. Heatmap of stable steady-state solutions for network shown in A, obtained via RACIPE. **C**. Density histogram of EM Score (= Zeb1 – miR-200) and Resistance Score (= ERα36 – ERα66) fitted to 3 and 2 gaussian distributions respectively. Red dotted lines show segregation between phenotypes: Epithelial (E), Hybrid (H) and Mesenchymal (M) for EM score, and Resistant (R) and Sensitive (S) phenotype for the latter. **D**. UMAP dimensionality reduction plots for steady-state solutions states obtained by RACIPE colored by either EM Score or Resistance score. **E**. Scatter plot showing corresponding EM Score and resistance score for all RACIPE solutions, and six biological phenotypes. Spearman correlation coefficient (ρ) and corresponding p-value are reported. **F**. Gene expression levels in six biologically defined phenotypes. Dotted line represents the monotonic increase in levels of ZEB1 across the phenotypes. Standard deviation is plotted as error bars. **G**. Density distribution of Spearman correlation coefficients (ρ) across an ensemble of 100 randomized versions of the GRN shown in A. Red line shows the correlation coefficient of the wild-type network shown in A.

To elucidate the emergent dynamics of this GRN, we simulated it using RACIPE (Random Circuit Perturbation) -a computational framework that solves a set of coupled ODEs (Ordinary differential equations) to examine the various phenotypic states enabled by the GRN, by sampling an ensemble of kinetic parameter sets from a biologically relevant parameter range (36). For each distinct parameter set, it then solves the ODEs to obtain possible steady states. For certain parameter sets, more than one steady state (phenotype) is achieved, suggesting possible stochastic switching among those states under the influence of biological noise.

Upon simulating our GRN using RACIPE, we observed multiple cell states that are visualized qualitatively as a hierarchically clustered heatmap (**Fig 1B**). Qualitatively, ZEB1, SLUG and ERα36 are often co-expressed and similarly, ERα66 and miR200 are co-expressed. The existence of these two major expression patterns is corroborated by K-means clustering (**Fig S1A**). Co-expression of ERα66 and miR200 can be construed as an Epithelial Sensitive (ES) phenotype, given that the presence of ERα66 associates with response to an anti-estrogen drug (e.g. Tamoxifen). However, in cases when ERα36 is higher, such cells are less likely to respond to an anti-estrogen compound and will exhibit drug tolerance or resistance. Thus, co-expression of ZEB1 and SLUG with ERα36 is interpreted as a Mesenchymal-Resistant (MR) phenotype. Further, we defined different cell states along the EMT and the drug resistance (TamR) axes. EM score is defined as the difference in normalized values of ZEB1 and miR-200; similarly, resistance score is defined as difference in normalized values of ERα36 and ERα66. Higher EM scores correspond to a mesenchymal phenotype, and higher resistance scores correspond to a resistant phenotype. A hierarchically clustered heatmap on the EM and resistance scores confirmed the existence of the ES and the MR phenotypes, along with the indication of other relatively less prevalent cell states such as Epithelial-Resistant (ER), Hybrid-Sensitive (HS) and Hybrid-Resistant (HR) phenotypes (**Fig S1B**).

Next, we plotted a normalized frequency histogram for the EM score of steady-state solutions obtained via RACIPE. The resultant distribution was visibly trimodal in nature (**Fig 1C**) with two dominant peaks corresponding to epithelial and mesenchymal phenotypes. The middle peak was smaller, suggesting the smaller abundance of hybrid E/M phenotypes expressing intermediate values of ZEB1 and miR-200, as earlier postulated (35) (**Fig S1C**). Similarly, the normalized frequency histogram of resistance score was bimodal, corroborated by existence of two distinct clusters along the ERα66-ERα36 axes (**Fig S1C**). To better visualize the various phenotypes enabled by the GRN, we performed UMAP analysis on the steady states and colored the individual steady states by either EM score or Resistance score. A qualitative comparison of the two UMAP plots revealed that the epithelial cluster was more likely to be sensitive while the mesenchymal cluster was more likely to be resistant (**Fig 1D**). The hybrid E/M (H) phenotype can be either Resistant (R) or Sensitive (S) (**Fig 1D, S1D**).

To further quantify the association between phenotypes on EM and resistance axes, we plotted the individual steady state solutions on these axes as a scatter plot. Discretization of the phenotypes along these axes (see **Methods**), and defined six possible distinct phenotypes – ES (Epithelial-Sensitive), ER (Epithelial-Resistant), HS (Hybrid-Sensitive), HR (Hybrid -Resistant), MS (Mesenchymal-Sensitive), and MR (Mesenchymal-Resistant) (**Fig 1E**). The ES and MR phenotypes were the most dominant while MS was the least prevalent. This analysis indicates that while there exists a strong association with the EM status and drug (tamoxifen) resistance at a cellular level (ρ = 0.806, p < 0.01), other phenotypes – ES, HR, HS and MS – may exist too, therefore highlighting the non-binary and semi-independent nature of EM phenotypes and its association with tamoxifen resistance.

Next, we assessed the molecular profiling of the six identified phenotypes. ZEB1 levels showed a monotonic increase in expression levels across the phenotypes ES, ER, HS, HR, MS and MR (**Fig 1F**). Intriguingly, SLUG levels, which was not used for classification of either EM or Resistance scores, appeared to be significantly higher in all the resistant states for a given EM phenotype, especially the hybrid E/M phenotype (**Fig 1F**). SLUG has previously been shown to be associated hybrid E/M states (35). Based on these observations, SLUG can be considered as a key player in driving resistance, especially in hybrid E/M phenotypes.

Finally, we examined whether this strong association between EM and Resistance scores/ phenotypes is specific to the network topology of this GRN only. To test this hypothesis, we created an ensemble of 100 randomized GRNs controlling for the total number of nodes in the network, net number of activation and inhibitory edges, and in and out degree of each node. We ran RACIPE simulations on this ensemble of networks and calculated the correlation coefficient between the EM and Resistance scores. The resulting distribution is centered around 0, and the ‘wild type’ GRN (**Fig 1A**) had the strongest correlation among the ensemble (**Fig 1G**), depicting the uniqueness of this network topology to this strong association. Further, we assessed the impact of other indirect gene regulatory links for ERα36, such as its self-activation and its ability to suppress the activity of ERα66 (16, 37). Addition of these links individually or in combination resulted in qualitatively similar results in the form of clustered heatmap and UMAP plots (**Fig S2**), suggesting that the GRN in **Fig 1A** is sufficient to capture fundamental features of EMP and reversible drug resistance observed in ER+ breast cancer.

### Clinical data support the predicted association between EMT and loss in ERα activity

As a preliminary validation of our model predictions, we probed the association between EMT programme and the loss in ERα activity with a concurrent gain in tamoxifen resistance markers. Specifically, we investigated clinical datasets of ER+ breast cancer patients treated with tamoxifen and observed that ESR1 (gene for ERα) levels were significantly negatively correlated with ZEB1 and SNAI2 (SLUG) levels, as well as with single-sample GSEA (ssGSEA) scores for MSigDB hallmark EMT signature (**Fig 2A**). Similarly, ssGSEA scores for signatures of resistance to tamoxifen at a proteomic level (TamRes) (38) were found to positively correlate with the levels of ZEB1, SNAI2 and MSigDB hallmark EMT programme (**Fig 2B**). Further, CDH1, a well-known epithelial marker (E-cadherin), was found to positively correlate with ESR1 levels, as well as with ssGSEA scores for MSigDB late estrogen response. Consistently, VIM, a canonical mesenchymal marker, showed negative correlation with the ssGSEA scores for MSigDB late estrogen response (**Fig 2C**).

**Figure 2:**
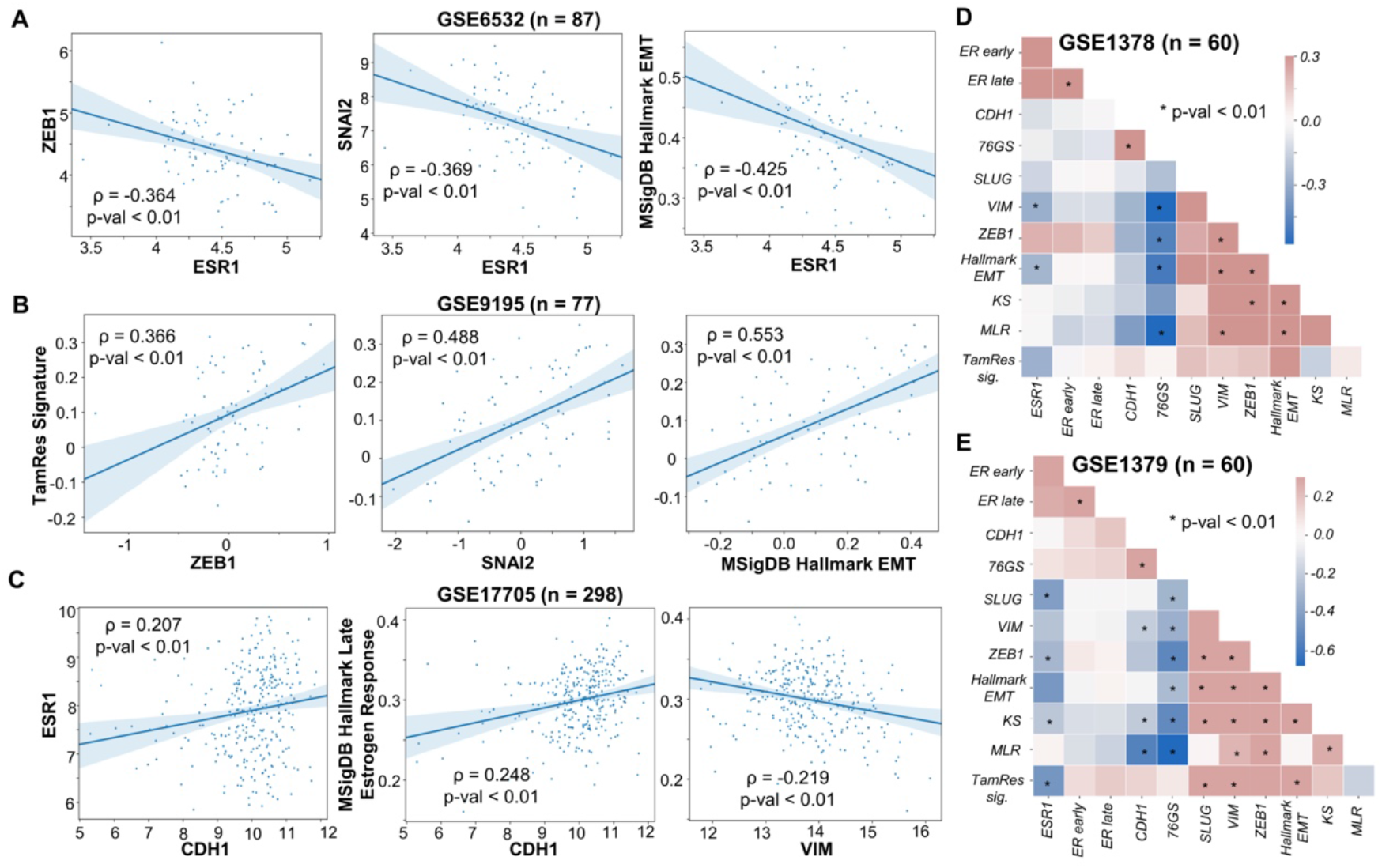
Gene expression analysis of publicly available datasets. **A**. Correlation of ESR1 with ZEB1, SNAI2 (SLUG) and activity of MSigDB Hallmark EMT signature (ssGSEA scores) in a cohort of 87 ER+ breast cancer patients (GSE6532). **B**. Correlation of Tamoxifen Resistance (ssGSEA scores) signature with expression of ZEB1 and SNAI2 and activity of Hallmark EMT signature in primary breast tumors treated with tamoxifen in adjuvant setting (GSE9195). **C**. Correlation of ESR1 expression levels and estrogen response activity with CDH1 and VIM in tumor samples from 298 ER+ patients treated with tamoxifen for 5 years. **D**. Diagonal correlation matrix between expression levels (ZEB1, SLUG, VIM, CDH1, ESR1), EMT scoring metrics (76GS, MLR and KS), gene set activity estimation (ER early response, ER late response, Tamoxifen resistance, Hallmark EMT signatures) for 60 samples of micro-dissected tumour biopsies (GSE1378) and whole tissue tumour biopsies (GSE1379) from a cohort of patients treated with Tamoxifen for 5 years. Spearman correlation coefficient (ρ) and corresponding p-value for scatter plots are reported.

Next, for a more comprehensive analysis of such correlations, we investigated pairwise correlations among expression levels of ESR1, canonical mesenchymal (SLUG, VIM, ZEB1) and epithelial (CDH1) genes, three EMT scoring metrics (76GS, KS, MLR) (39), and four gene set activity estimation via ssGSEA scores (ERα early response, ERα late response, Tamoxifen resistance and Hallmark EMT). Across patient samples irrespective of whether the samples were micro-dissected tumour biopsies or whole tissue tumour biopsies, we observed an expected positive correlation among EMT metrics, mesenchymal markers and ssGSEA scores for hallmark EMT (**Fig 2D**). ESR1 gene expression levels usually correlated negatively with mesenchymal markers and/or EMT scoring metrics. Conversely, Tamoxifen resistance signature correlated positively with the mesenchymal markers and EMT ssGSEA scores, but negatively with ESR1 (**Fig 2D**). Put together, this analysis supports our prediction about an association between activation of EMT programme and compromised ERα signaling activity.

### Reciprocal driving of the EMT programme by suppressing Estrogen receptor activity and *vice versa*

After investigating the correlations among EMT and tamoxifen resistance axes, we inspected whether these processes could drive one another. To understand the effects of perturbations of the EM axis on estrogen signaling axis and drug resistance, we first simulated overexpression (OE) and downexpression (DE) of ZEB1. Over expression of ZEB1 led to a significant increase in the frequency of MR phenotype with concurrent decrease in ES, HR and ER phenotypes (**Fig 3A**). Opposite trends were seen on in ZEB1 DE case, as expected. An increase in MR phenotype upon ZEB1 over-expression indicates that as cells are driven to undergo EMT via ZEB1, they lose their sensitivity to anti-estrogen drugs. This change should reflect as a decrease in estrogen signaling activity in cells. To test this hypothesis, we analyzed publicly available gene expression data for ER+ breast cells/cell lines induced to undergo EMT. In HMLE cells where EMT was induced by over-expression of Twist, Snail or Slug, ssGSEA scores for Hallmark EMT gene list showed significant increase in enrichment levels in the EMT program with a concurrent decrease in the activity of gene lists representing early Estrogen Receptor response and late Estrogen Receptor response (**Fig 3B**). Similarly, in other datasets, EMT induction via TGFβ, Twist, Gsc or Snail in HMLE cells, or that via Six1 over-expression in MCF7 (ER+ breast cancer cell line) cells showed reinforcing trends including enrichment of the Tamoxifen Resistance program (**Fig 3C, S3A**). Over-expression of SNAIL in MCF10A upregulated ZEB1 and the EMT program and had downregulation of CDH1 levels as well as a concurrent drop in Estrogen Receptor activity (both early response and late response) (**Fig S3B**). Interestingly, SLUG levels were reduced upon SNAIL over-expression, reminiscent of reports about mutual repression between SNAIL and SLUG (40). Given that SLUG can induce a partial EMT while SNAIL is likely to induce more of a complete EMT (35, 41), these results together suggest that drug resistance can be achieved even through a partial EMT state, and that SNAIL and SLUG may follow different paths in the multi-dimensional EMT landscape, both of which can confer tamoxifen resistance to ER+ breast cancer cells.

**Figure 3:**
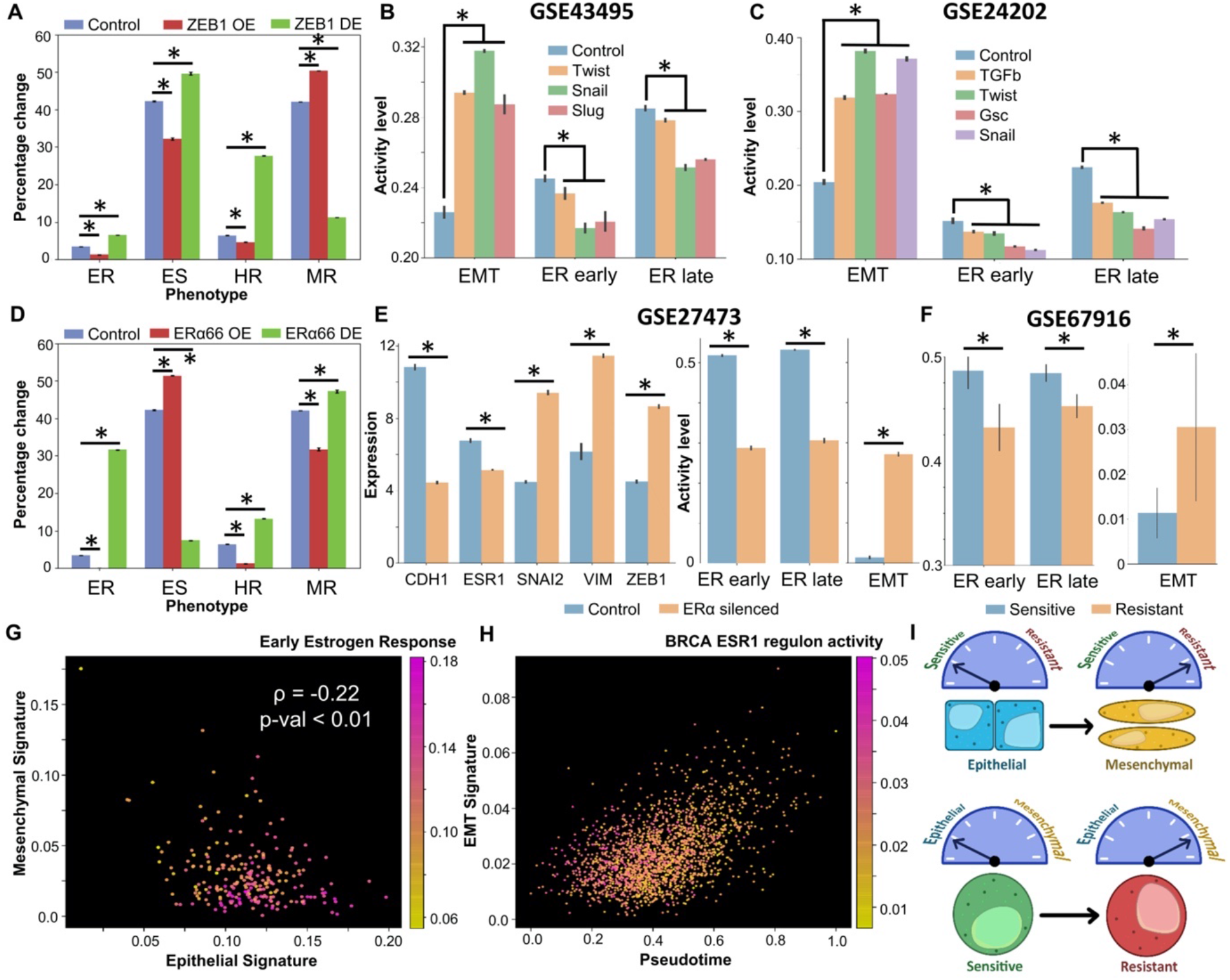
Induction of EMT can drive suppression of Estrogen signaling and vice versa. **A**. Impact of over-expression/down-expression of ZEB1 levels in RACIPE simulations on frequencies of different biological phenotypes. Error bars denote standard deviation across n=3 replicates. **B**. Experimental data (GSE43495) for EMT induction via Twist, Snail or Slug in HMLE cells and the concurrent decrease in the magnitude of early and late estrogen response. **C**. Experimental data showing EMT induction via TGFβ, Twist, Gsc and Snail in HMLE epithelial cells and the concurrent decrease in the magnitude of early and late estrogen response (GSE24202). **D**. Same as A. but **for** over-expression/down-expression of ERα66. **E**. Experimental data showing differences in gene expression levels of Cdh1, Vim, Snai2 (Slug), Vim and Zeb1 and change in magnitude of early and late estrogen response and the EMT programme (ssGSEA on MSigDB Hallmark EMT signature) in control and ERα silenced MCF7 cells (GSE27473). **F**. Experimental data showing differences in activity levels of early ER response, late ER response and EMT programme in sensitive and resistant MCF7 cell lines (GSE67916). For A-F, * denotes a statistically significant difference between the control and perturbed/induced case assessed by a two-tailed Students t-test assuming unequal variances. **G**. Scatter plot showing association between activity of early estrogen response and cells with varying positions on a 2D epithelial-mesenchymal plane (GSE147356). Spearman’s correlation coefficient between epithelial and mesenchymal scores, and corresponding p-value are reported. **H**. Scatter plot showing activity of EMT signature in TGFβ treated MCF7 individual cells in pseudo time and the concurrent decrease in BRCA ESR1 regulon activity. Color bar represents the range of activity level of the ESR1 regulon (GSE147405). **I**. Schematic showing bidirectional associations between the EMP and the drug resistance programme, i.e. induction of EMT drives a switch to a therapy-resistant state, and acquisition of therapy resistance often drives EMT.

Next, we inquired whether perturbing the levels of ERα66 could lead to a more mesenchymal phenotype by inducing either a partial or complete EMT. We simulated the GRN by RACIPE with both over-expression (OE) and down-expression (DE) of ERα66, and observed that the over-expression of ERα66 caused a marked reduction in the levels of all resistant phenotypes with a concomitant increase in epithelial sensitive (ES) phenotypes (**Fig 3D**). Conversely, downregulation of ERα66 gene caused a significant drop in the prevalence of ES phenotype with cells being pushed towards an ER, HR or MR phenotype (**Fig 3D**). Thus, the impact of ERα66 down-regulation can be multi-faceted where it could drive the system towards a more resistant state without changing the EMT status (i.e. to ER phenotype) or could push the cells to a resistant state by making cells more mesenchymal. Interestingly, when ERα was silenced in MCF7 cells, there was a marked increase in the levels mesenchymal genes such as SLUG, VIM and ZEB1 with a significant decrease the levels of CDH1 (**Fig 3E**). The effect of ERα silencing was also observed in decreased early and late Estrogen receptor activities, and a concurrent increase in the activity levels of the EMT hallmark gene signature (**Fig 3E**). This result establishes that ERα silencing alone can drive EMT and consequently exhibit a drug resistant phenotype. Further, we observed that in resistant MCF7 cells derived from the sensitive parental population, the estrogen response pathways were significantly suppressed, together with upregulation of EMT program (**Fig 3F**). In another dataset consisting of sensitive and resistant cells, both CDH1 and ESR1 levels were significantly lower in resistant cells and SLUG was significantly upregulated (**Fig S3C**). Although ZEB1 and VIM were not significantly upregulated, the overall EMT program was higher in resistant cells than in sensitive breast cancer cells (**Fig S3C**). Given the profiles of CDH1, ZEB1, ESR1 and VIM in these MCF7 cells, they could be construed as a hybrid-resistant (HR) phenotype, at least at a bulk level.

Further, we evaluated whether these trends were also preserved at a single cell level. We first investigated the 10 cell sequencing data of ER+ breast cancer cells (42). The 2D scatter of activity levels of epithelial and mesenchymal genes revealed an expected negative correlation denoting the reciprocal epithelial and mesenchymal phenotypes (**Fig 3G**). Intriguingly, we found that epithelial cells were more likely to harbor a responsive estrogen receptor pathway (**Fig 3G**). Further, the activities of early and late estrogen receptor hallmark genes showed a strong positive correlation with the 76GS EMT scoring method where higher scores indicate a more epithelial phenotype (**Fig S3D**). The activity of BRCA specific ESR1 regulon was also found to be significantly positively correlated with 76GS EMT scoring metric, suggestive of an active estrogen receptor signaling in epithelial cells rather than mesenchymal ones (**Fig S3D**). Finally, we examined if EMT induction at a single cell level could itself drive a suppression of the estrogen receptor pathway activity. In MCF7 cells treated with TGFβ to induce EMT, we plotted the EMT program activity as a function of pseudo-time as reported earlier (43). EMT activity score showed a significant positive correlation with the pseudo-time, establishing pseudo-time as a proxy for EMT progression. Upon coloring by the BRCA specific ESR1 regulon activity, we found that lower values of EMT activity were more likely to be associated with a higher ESR1 regulon activity level (**Fig 3H**). We confirmed this observation by plotting the activities of early and the late estrogen receptor hallmark genes and the ESR1 regulon with pseudo-time. All of them showed a significant negative correlation with pseudo-time, further supporting the earlier observation (**Fig S3E**). Overall, we demonstrated that induction of EMT can suppress estrogen signaling axis and vice versa, resulting in concurrent change in cellular phenotypes along both the EMP axis and in levels of drug resistance in ER+ breast cancer cells (**Fig 3I**).

### Stochastic transitions between different phenotypes on EMT and drug resistance axes

Next, we investigated whether under the influence of biological noise (44), these different phenotypes can switch among one another on EMT and/or tamoxifen resistance axes. We identified the parameter sets simulated via RACIPE that gave rise to multi-stability (i.e. more than one steady state solutions, depending on the initial condition chosen) and performed stochastic simulations.

We first characterized the proportion of parameter sets simulated by RACIPE that are mono-stable vs multi-stable. Approximately two-third (∼65%) of the parameter sets were found to be bistable, followed by 20% parameter sets exhibiting tristability, and 8% monostable cases (**Fig 4A**), indicating that the GRN simulated is poised for multistability. Within the monostable solutions obtained, {ES} and {MR} are the most predominant phases. Among bistable cases, the most common resultant phase is that of {ES, MR}, followed by {HS, MR} and by {ES, HR} Finally, among tristable cases, {ES, HR, MR} phase is the most predominant followed by {ES, ER, MR} and then by {ES, HS, MR} (**Fig 4A**). Put together, these results suggest that epithelial sensitive and mesenchymal resistant are likely to be most frequent subpopulations in a given cancer cell populations, with smaller proportions of hybrid E/M cells which may be tamoxifen-sensitive or tamoxifen-resistant.

**Figure 4:**
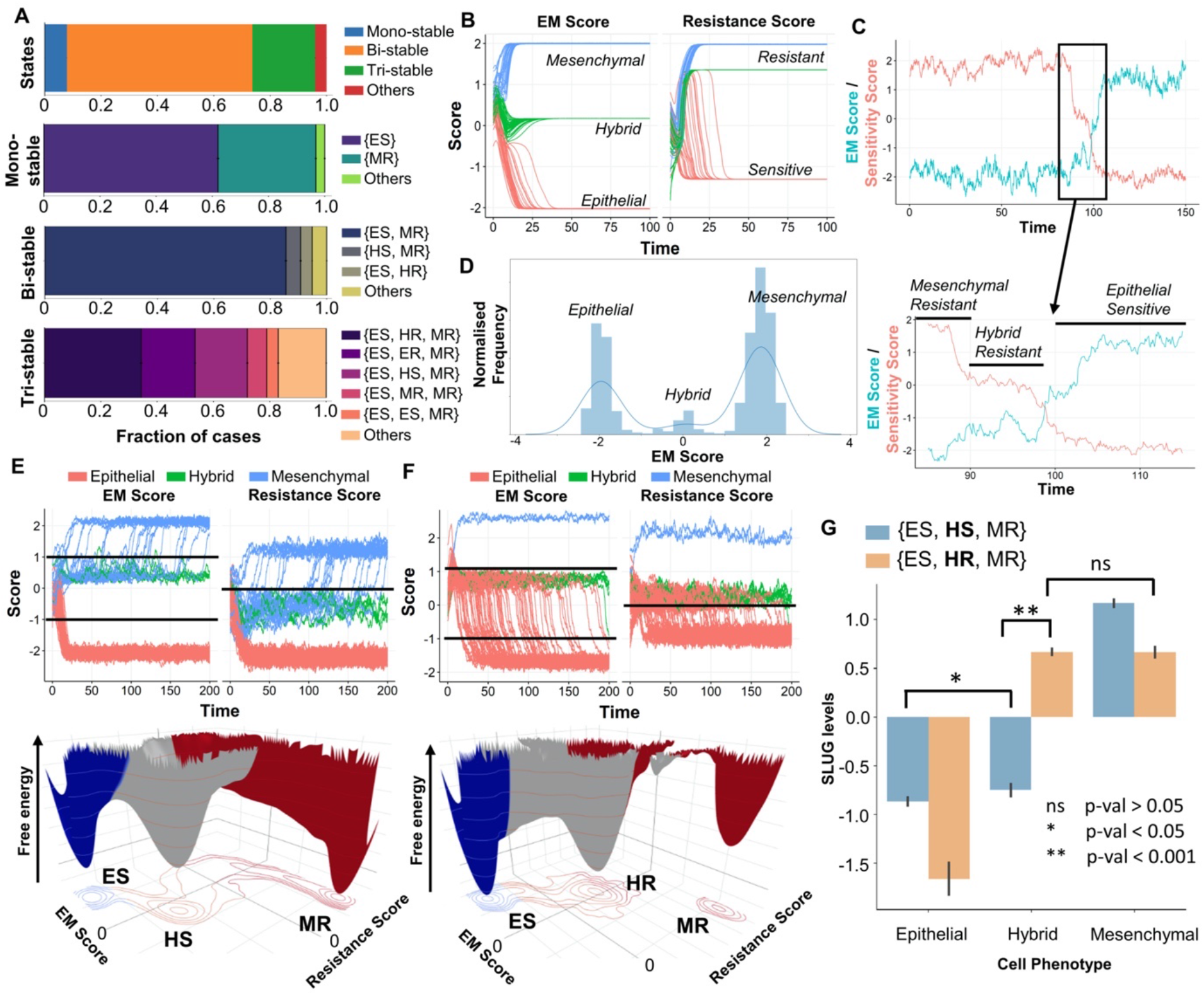
Stochastic stimulations showing dynamic state transitions among different biological phenotypes. **A**. Fraction of RACIPE parameter sets resulting in monostable and multi-stable solutions (bi-, tri-, others) and the frequency distribution of phases that compose the monostable and multi-stable solution sets. **B**. System dynamics for a representative {ES, HR, MR} parameter set showing the existence of the 3 biological EM phenotypes (E, H, M) and resistant (R) and sensitive (S) phenotypes when started from multiple initial conditions. **C**. Time course showing the transition of the system from a MR to an ES phenotype through a HR state under the influence of noise. Sensitivity score is defined as negative of the tamoxifen resistance score, i.e. ERα66 – ERα36. **D**. Marginal distribution of the EM score from the time course shown in C; 3 peaks denote existence of 3 distinct states along EM spectrum. **E**. (top) Stochastic time series for multiple initial conditions tracking EM and Resistance scores in a representative parameter set from the {ES, HS, MR} phase. (bottom) Landscape obtained by simulation of that parameter set with valleys representing stable states possible in the system. **F**. Same as E. but for a representative parameter set from the {ES, HR, MR} phase. **G**. Changes in SLUG levels as the system transitions from ES to MR phenotype through either HS or HR state. HR state is characterized by high levels of SLUG compared to HS cell state. Students’ t-test results show the level of statistical significance between various comparisons.

We focused on tristable parameter sets that contain ES and MR phenotypes and can enable transition between them through an intermediate state. We considered a representative parameter set from the {ES, HR, MR} phase, and plotted the levels of EM Score (ZEB1 – miR200 levels on log2 scale) and the Resistance score (ERα36 – ERα66 levels on log2 scale), sampling multiple possible initial conditions. As expected, we observed three distinct levels in steady-state EM scores, corresponding to one each in the E, H or M region. The resistance score also showed three distinct levels; however, two of them were classified as R phenotype with one of them as S (**Fig 4B**). Stochastic simulations for this parameter set revealed noise-induced switching from a mesenchymal-resistant (MR) phenotype to an epithelial-sensitive (ES) phenotype through a hybrid-resistant (HR) phenotype (**Fig 4C**). The existence of these three states was further corroborated by plotting the marginal distribution of the EM Score obtained from the time profile, that revealed three peaks with varying EMT scores (**Fig 4D**).

Next, we constructed landscapes to interpret cell-state transitions possible in the co-existing phenotypic combinations of {ES, HS, MR} and {ES, HR, MR}. To do so, we simulated the system from multiple initial conditions under the influence of noise, and obtained the pseudo potential of the points in state-space as the negative log of the probability of occurrence. We observed that for the representative case from the phase {ES, HS, MR}, trajectories that started out as hybrid-sensitive phenotype (HS) switched to a mesenchymal-resistant (MR) phenotype (**Fig 4E**, top), thus unraveling the dynamic nature of the observed cell states. The landscape constructed for this parameter set revealed 3 distinct valleys or “attractors” – ES, HS and MR (**Fig 4E**, bottom). For a representative parameter set from the phase {ES, HR, MR}, we again saw 3 distinct EM states and the corresponding expected resistant states (**Fig 4F**, top). The three “attractors” observed here was shifted along the axis of the resistance score – from < 0 for HS to > 0 to HR (**Fig 4F**, bottom vs. **Fig 4E**, bottom). Similar dynamics were observed for other kinetic parameter sets belonging to these two tristable phases (**Fig S4A, B**). A key difference noted in the HS and HR states was the levels of SLUG, suggesting SLUG as a potential marker for hybrid E/M resistant phenotype (**Fig 4G**).

These results demonstrate that the different states in the two-dimensional space of EMT and tamoxifen resistance (i.e. non-genetic heterogeneity) can also switch among one another (i.e. phenotypic plasticity) under stochastic variations in gene expression and/or biochemical rates. It should be noted that these state transitions can also be driven through external perturbations such as treatment with TGFβ or with anti-estrogen treatments such as tamoxifen. Simulations shown here demonstrate the paths cells traverse through while transitioning to other state(s).

### Complementary roles of heterogeneity and plasticity for the tumour survival of ER+ breast cancer cells under anti-estrogen treatments

After elucidating the emergent intra-cellular dynamics of the coupled gene regulatory network (GRN) of EMT and ERα signaling, we probed the effect of the dynamical traits of this network – phenotypic plasticity and non-genetic heterogeneity – at a cell population level. To gain a better understanding of how these two cell-autonomous traits (a) heterogeneity in sensitivity of a cell towards a drug, and b) ability of cells to switch bidirectionally between a sensitive and a resistant phenotype) can influence the long-term survival of a cell population, we developed a simple mathematical model involving only 2 components – drug sensitive (S) cells and drug resistant (R) cells. Although non-cell autonomous phenomenon such as competition and/or cooperation between sensitive and resistant cells (45–47) have been extensively observed and mathematically modelled, it remains unclear how plasticity and/or heterogeneity in cell populations might contribute to long term population survival, even in absence of such non-cell autonomous behaviors.

In this modeling framework, cells can belong to either a sensitive (S) phenotype or a resistant (R) phenotype. The degree of sensitivity of a cell to a drug is defined by a “resistance score” which is sampled from a gaussian distribution with a given mean and standard deviation. Heterogeneity in the system is modelled via changing the standard deviation of the gaussians from which the resistance score is sampled. The sensitivity of each cell (probability that the cell escapes drug-induced killing) depends on the resistance score through a sigmoidal curve (**Fig 5A**), reminiscent of typical IC50 curves. Further, the cells can switch from a sensitive to a resistant phenotype with a given probability P_SR_ and can switch back with a probability P_RS_, thus introducing plasticity into the system (**Fig 5A**). These probabilities can depend on various external conditions such as drug exposure time and/or drug concentration.

**Figure 5:**
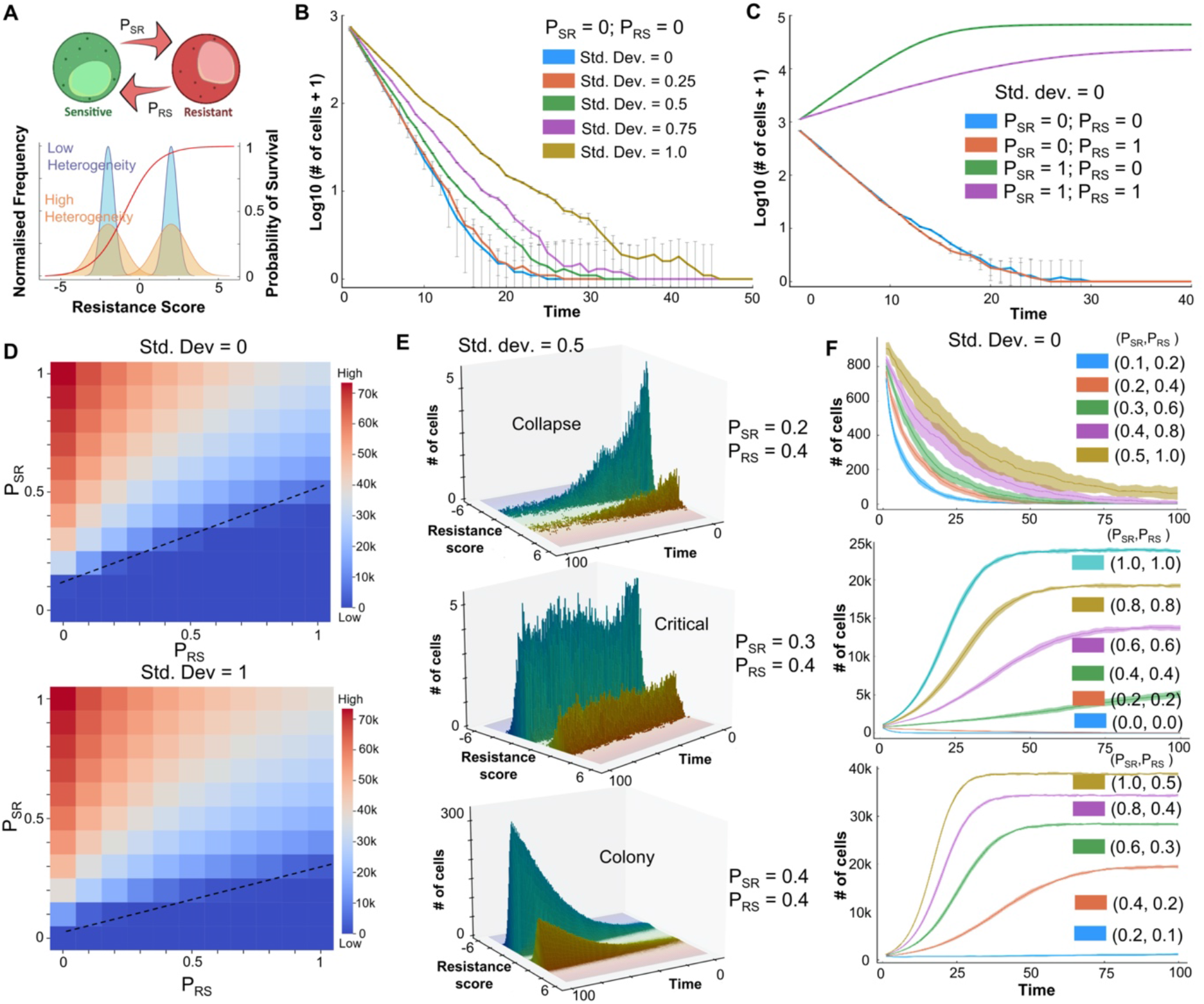
Effect of heterogeneity and plasticity on tumor survival in the presence of an anti-estrogen drug. **A**. Schematic for model formulation showing inter-conversions between sensitive and resistant phenotypes (transition probabilities: P_SR_, P_RS_). Heterogeneity in the cell population is modelled by standard deviation (Std. dev.) of Gaussians from which resistance scores are sampled. Survival probability of cells is a function of resistance score approximated as a sigmoidal curve (shown in red). **B**. Effect of heterogeneity on population sizes over time, starting with an initially all sensitive cell population and at P_SR_ = P_RS_ = 0. **C**. Effect of plasticity on population sizes over time starting with an initially all sensitive cell population and no heterogeneity (Std. dev. = 0). **D**. Population sizes (at time(t) = 100) as a function of P_SR_ and P_RS_ at two different heterogeneity levels, starting with an initially all sensitive cell population. Dotted lines indicate a qualitative boundary between tumor survival and elimination scenarios. **E**. Distinct qualitative scenarios – collapse of initial population of cells, maintenance of the cell population around starting initial conditions (in the time frame considered) and net growth in a population of cells leading to survival of the tumour – at varying levels of P_SR_ and P_RS_. All simulations start with a fixed heterogeneity (Std. dev. = 0.5) and a fully sensitive population. **F**. Population sizes over time as a function of varying values of (P_SR_, P_RS_). All simulations start with no heterogeneity (Std. dev. = 0) and a fully sensitive population.

Simulations for this model showed that in the absence of any drug, an initially fully sensitive population of cells with no heterogeneity (resistance score for all cells < 0, std. dev. = 0) and no plasticity (P_SR_ = P_RS_ =0) could grow to and eventually saturate to a population size close to the carrying capacity (10^5^ cells) (**Fig S5A**; blue curve). However, the presence of drug can eliminate this population of non-plastic and homogeneous drug-sensitive cells (**Fig S5A**; green curve); the rate of elimination depends on intrinsic growth and death rates of cells (**Fig S5A**; orange and violet curves). Next, we characterized the role of increasing heterogeneity in a system devoid of plasticity (P_SR_ = P_RS_ =0). We observed that increasing heterogeneity (shown by different standard deviation values) can delay the time taken to eliminate a population of cells, but it was not sufficient in enabling the survival of population, as long as the population overall is predominantly sensitive as per the survival probability curve (**Fig 5B, S5B-E**).

Next, we examined the influence of plasticity in the absence of heterogeneity (std. dev. = 0). For P_SR_ = 0 (no switching from a sensitive to a resistant phenotype), irrespective of the value of P_RS_, the cell population is eliminated (**Fig 5C**; blue and orange curves). For P_RS_ = 0 (no switching from resistant to sensitive population), if the probability of switching from sensitive to resistant is very high (P_SR_ = 1), the population of cells survive. However, increasing values of P_RS_ can decrease the final population size (**Fig 5C;** green and purple profiles). Further, we performed an exhaustive analysis of the P_SR_ -P_RS_ plane with values ranging from 0 to 1 with steps of 0.1 and colored the matrix based on the final population size at time t = 100 steps. We first performed this analysis for two extreme values of heterogeneity (std. dev. = 0 and 1). We observed that in presence of higher heterogeneity, population survival and growth was seen for more combinations of (P_SR_, P_RS_) values (compare **Fig 5D**; top and bottom panels). Further, if P_SR_ << P_RS_, an increase in heterogeneity alone cannot rescue the population and the population is eliminated. On the other hand, if P_SR_ >> P_RS_, the population survives and reaches near maximum colony sizes (close to carrying capacity in the system) (**Fig 5D**). These trends were qualitatively similar for other levels of heterogeneity (**Fig S6**) as well, highlighting potentially universal organizing principles in determining cancer cell population fitness.

Finally, we characterized different qualitative properties exhibited by a cell population under varying values of P_SR_ and P_RS_, but at a fixed heterogeneity level. To visualize the proportion of cells in the sensitive and the resistant component separately, we plotted a 3D histogram showing the temporal evolution of cell population, colored by corresponding resistance scores. For an intermediate value of heterogeneity, depending on the relative values of P_SR_ and P_RS_, three distinct qualitative outcomes for cell population are possible: a) it can be eliminated completely, b) it can maintain its critical population size similar in magnitude to the initial number of cells, c) it can increase rapidly to saturate at a higher value to establish a colony (**Fig 5E**). Next we explored the effect of relative and absolute values of P_SR_ and P_RS_ on the overall population dynamics of the system. In absence of heterogeneity, as a representative case, we simulated various scenarios where the ratio P_SR_/P_RS_ was fixed to be 0.5, 1 or 2. When P_SR_/P_RS_ = 0.5, the population always collapsed irrespective of absolute values of P_SR_ and P_RS_ (**Fig 5F**). However simulation cases that had higher absolute values of P_SR_ and P_RS_ showed a slower rate of extinction; this trend was seen also for P_SR_/P_RS_ = 1 and 2 (**Fig 5F**). These cases highlight that not only the relative rates of plasticity in either direction (S to R or vice-versa), but also absolute residence times (proxied by P_SR_ and P_RS_) can influence population dynamics, by modulating the time of exposure of cells to the therapy. As expected, higher P_SR_/P_RS_ increases the propensity of population survival, and thus a faster growth curve and higher colony size. Collectively, these results indicate the complementary roles of heterogeneity and plasticity in the cell population to determine the survival probability of a population of cancer cells in the presence of an anti-estrogen therapy such as tamoxifen.

### “What does not kill [cancer cells] makes them stronger” – the influence of drug-induced plasticity and intrinsic non-genetic heterogeneity in population survival

As described above, depending on the absolute values of P_SR_ and P_RS_, a population of cells can collapse, grow a colony or maintain the population size around the initial value. The last case is likely to be a metastable state as under the effect of intrinsic noise, the population can be pushed to be eliminated completely or grow and saturate to carrying capacity of the system. This feature enables us to define the extinction probability for a cell population, for a set of simulations from specified initial conditions. Extinction probability is defined as the fraction of cases in which the population is eliminated after a long time. For most values of P_SR_ and P_RS_ the extinction probability is either 0 (population always grows and establishes a fixed colony) or is 1 (population is eliminated), under the influence of no heterogeneity (std. dev. =0). We explored whether increasing heterogeneity can have an extinction probability between 0 and 1. As a representative case, we found that starting from an initially all sensitive population (resistance score < 0), with P_SR_ = 0.5 and P_RS_ = 1, intermediate heterogeneity level (std. dev. = 0.65) led to extinction of the population (**Fig 6A**, left; extinction probability = 1.0). However, increasing the heterogeneity (std. dev. = 0.7) led to a decrease in extinction probability (0.43 +/-0.03) with the surviving cases saturating at a population level of ∼750 cells (**Fig 6A**, middle). A further increase in heterogeneity further reduced extinction probability (0.14 +/-0.02) concomitant with an increasing saturating population size of ∼1500 cells (**Fig 6A**, right). These simulations illustrate a protective role of non-genetic heterogeneity in maintaining a cancer cell population, under the influence of therapy, especially at high enough intrinsic switching rate of cells to a resistant state, due to stochastic dynamics of phenomenon such as EMT.

**Figure 6:**
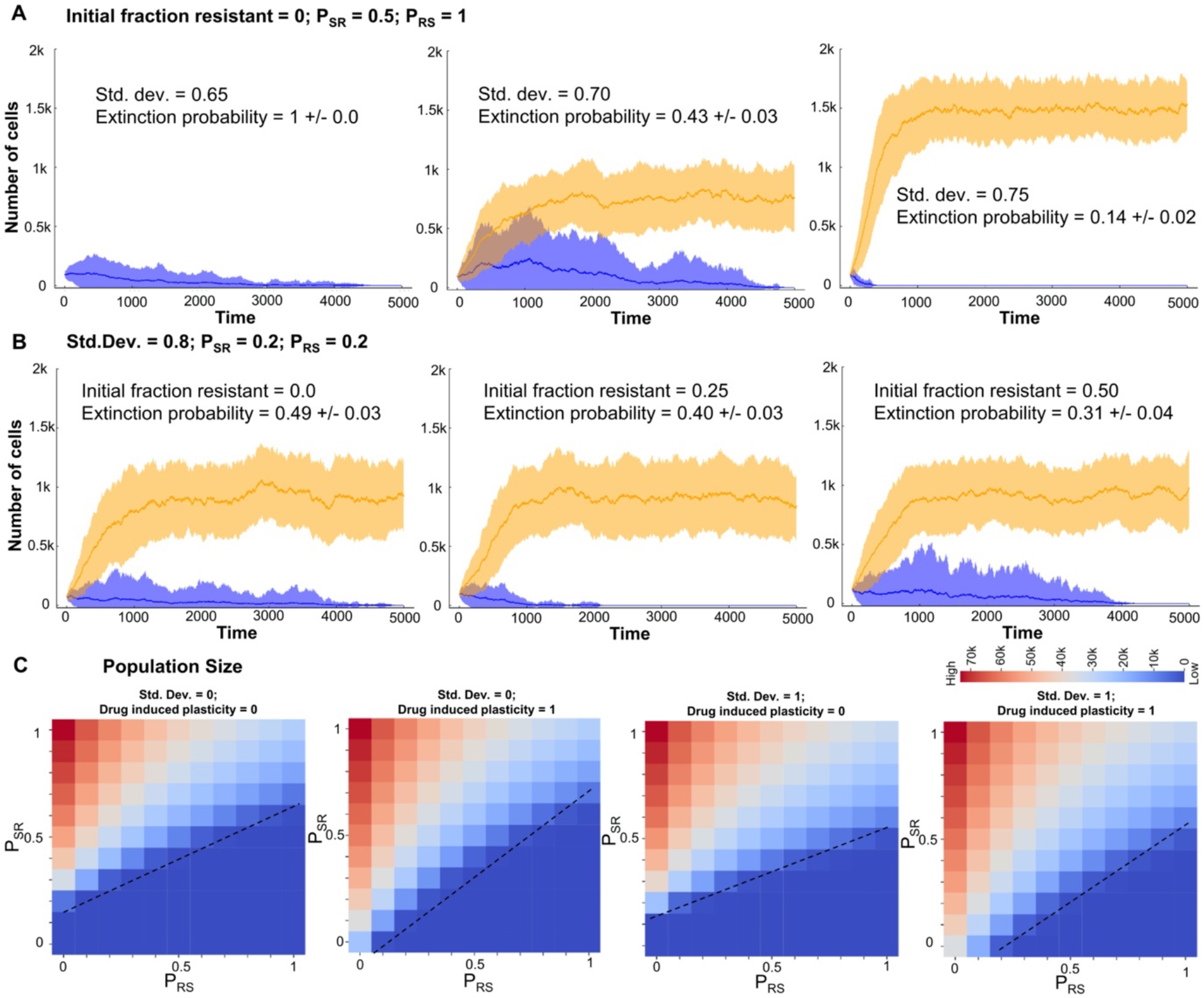
Different mechanisms promoting survival of a cancer cell population in the presence of an anti-estrogen drug. **A**. Simulations showing increase in survival probability (decrease in extinction probability) of a cancer cell population with varying heterogeneity levels (Std. dev. = 0.65, 0.70 and 0.75) starting form an all sensitive population and fixed values of P_SR_ and P_RS_ at 0.5 and 1.0 respectively. **B**. Simulations showing a modest increase in the survival probability of a cell population with varying levels of initially resistant cells (initial fraction = 0.0, 0.25 and 0.50) and fixed values of P_SR_ and P_RS_ at 0.5 and 1.0 respectively. Orange ribbon represents the collection of all states that survive and form a colony and blue ribbon for all those cases that are eventually eliminated. The dark line represents the mean and the band represents the std. dev. around that mean for an ensemble of simulations. **C**. Final population sizes (at time(t) = 100) as a function of P_SR_ & P_RS_ at two different heterogeneity levels (Std. dev. = 0, 1) and two different levels of drug induced plasticity (0 and 1) starting with an initially all sensitive cell population. Dotted lines indicate a qualitative boundary between cases where a tumour survives or is eliminated by the presence of the drug.

Next, we investigated the effect of initial fraction of “pre-existing resistant cells” (48) on the population extinction probability. At fixed heterogeneity (std. dev. = 0.8) and plasticity levels (P_SR_ = P_RS_ = 0.2), starting with only sensitive cells (resistance score < 0), the extinction probability was 0.49 +/-0.03, but increasing the initial fraction to 0.25 (i.e. 25% of initial cells have resistance score > 0) caused a modest decrease in extinction probability (0.40 +/-0.03) (**Fig6B**), as compared to the impact of heterogeneity, where a decrease of approximately 60% was noted for extinction probability (**Fig 6A**). A further increase in this initial fraction also had a similar weak effect in changing either the extinction probability (0.31 +/-0.04) or the final population size (**Fig 6B**). This analysis revealed that the initial fraction of reversibly resistant cells is perhaps a weaker factor, especially at lower intrinsic transition probabilities between sensitive and resistant cells.

Finally, we investigated the effect of drug-induced plasticity (i.e. externally induced transitions from a sensitive to a resistant state) (9) on the cell population dynamics. In our model, sensitive cells that can survive drug exposure in stipulated time have a probability to switch to resistant state. We monitored the final population size with varying levels of drug induced plasticity (0 and 1) at two different levels of heterogeneity (std. dev. = 0 or 1). We observed that under high drug-induced plasticity conditions, especially with smaller P_RS_ values, a marked increase in final population size was observed, irrespective of the extent of heterogeneity (**Fig 6C, S7**).

Put together, these observations show diverse mechanisms that can promote the survival of a cancer cell population under the influence of various anti-estrogen drugs: a) non-genetic heterogeneity in initial cell population, including the percentage of *de novo* reversibly resistant cells, and b) stochastic switching between cell states (ES, MR) under the influence of noise, maintaining a dynamic equilibrium of sensitive and resistant cells, and c) drug-induced switch to a reversible resistant state for cells that do not die upon exposure of the drug, i.e. cells that tend to follow the Nietzsche’s proposal of “what does not kill me makes me stronger” (49).

### Combinatorial strategies to target a plastic and heterogeneous cancer cell population

After deciphering multiple possible routes to long-term sustenance of a resistant population which can lead to tumor growth, we investigated what mechanisms can be efficiently deployed to target a plastic and heterogeneous cancer cell population. From our network simulations and follow-up analysis of bulk and single-cell transcriptomic data, we established a consistent association between epithelial state and higher drug sensitivity, as well as a mesenchymal one exhibiting recalcitrance to tamoxifen (**Fig 1, 2**). Thus, in our cell population dynamics framework, we incorporated the effect of an MET-inducer, i.e. an external signal that increased P_RS,_ just as drug-induced switching increased P_SR_. Thus, this MET-induced sensitivity forced cells to switch from a more mesenchymal (i.e. resistant) to an epithelial (i.e. sensitive) cells, thus reducing the fraction of resistant cells in the population.

To simulate the effects of such a scenario, we first chose a representative parameter set that enabled a surviving population of cells, with a larger frequency of sensitive cells than resistant ones (**Fig 7A, i**). Upon introducing drug induced plasticity into the system, we observed faster saturation, and a changed demographic with most of the population consisting of resistant cells (**Fig 7A, ii**). Upon introduction of the MET inducer in the absence of drug induced plasticity, the population undergoes a rapid collapse as the cells are sensitized to the drug, causing drugs to kill the cells (**Fig 7A, iii**). However, in the presence of drug induced plasticity, the population undergoes a relatively slower collapse as these two factors (drug-induced plasticity and MET-induced sensitivity) are opposing in nature (**Fig 7A, iv**).

**Figure 7:**
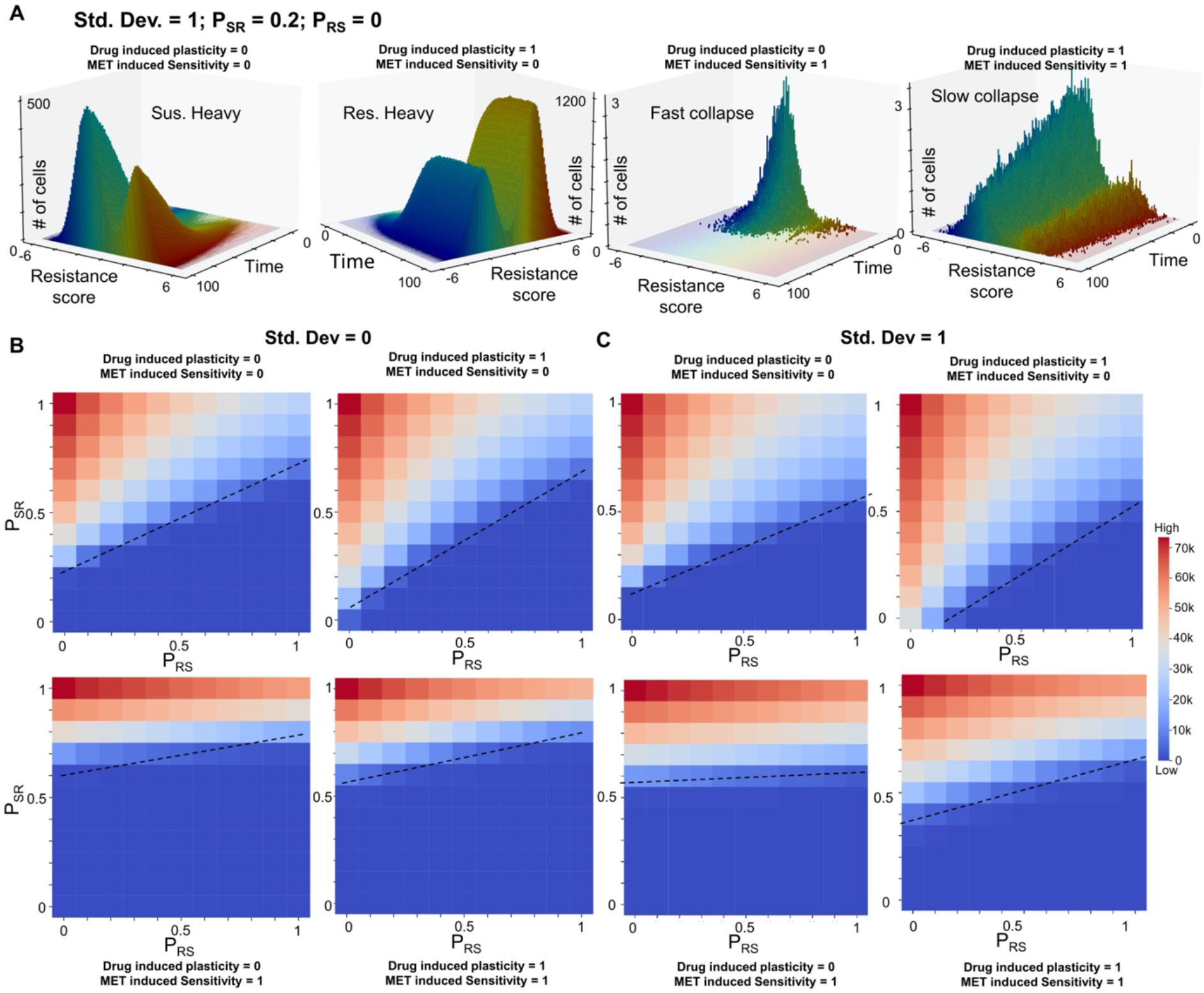
MET inducer, in conjunction with anti-estrogen drugs, can potentially limit the survival of cancer cell population. **A**. Temporal evolution of a population of cancer cells (resistant and sensitive both) under different levels of drug induced plasticity (0 and 1) and MET induced sensitivity (0 and 1). Simulations were performed starting form an all sensitive population and fixed values of P_SR_ and P_RS_ at 0.2 and 0 respectively. **B**. Final population sizes (at time(t) = 100) as a function of P_SR_ and P_RS_ 2 different levels of drug induced plasticity (0.1 and 1) and 2 different levels of MET induced drug sensitivity (0 and 1) starting with an initially all sensitive cell population with no heterogeneity in the system. Dotted lines indicate a qualitative boundary between cases where a tumour survives or is eliminated by the presence of the drug.

This behaviour is generally conserved across multiple parameter values corresponding to heterogeneity and plasticity (std. dev., P_SR_, P_RS_). Irrespective of the extent of heterogeneity, we see a larger combination of (P_SR_, P_RS_) values in the P_SR_ -P_RS_ plane allowing for population collapse in the presence of MET-induced sensitivity as compared to the control case (**Fig 7B**). These observations suggest that strategies that can revert the influence of therapy-induced plasticity to alter the population demographic to higher proportion of sensitive cells can be applied together with canonical anti-estrogen therapies to halt the dynamic adaptability of cell population, thus “trapping” them in a drug-sensitive state. In the current paradigm, ‘tolerant’ cells which are able to escape being killed upon drug exposure are very likely to switch to a resistant state, given the coupled EMT-estrogen signaling coupling, thus adding to existing tumor burden. However, alleviating this side-effect of the drug-induced plasticity through MET inducers which may sensitize the population, can be a more efficient therapeutic combination.

## Discussion

Drug resistance, similar to many hallmarks of cancer, has traditionally been presumed to have a strong genetic underpinning. Thus, drug-resistant cells in a tumor population have been thought of containing pre-existing (*de novo*) mutations or in acquired mutations during the course of therapy (2). However, over the past decade, accumulating experimental/preclinical and clinical evidence of non-genetic, stochastic and reversible modes of drug resistance – labelled often as Drug-tolerant persisters (DTPs) – have been reported *in vitro* and *in vivo* (5, 50–53), reminiscent of similar observations in microbial systems (54). DTPs have been also shown to give rise to stable drug-resistant phenotypes involving genetic changes upon prolonged growth (53), highlighting the importance of realizing different timescales over which cellular adaptation can occur. While short-term changes are often phenotypic in nature and involve foraging of the phenotypic landscape, long-term survival strategies may involve genomic changes to enable being trapped in the “attractors” explored during short-term foraging. While such transitions to a persister (reversibly resistant) cell state have been largely associated with epigenetic reprogramming (2), recent studies show significant variation in transcriptional programme of such cells too as compared to sensitive cells from the same genetic background (48, 55–57), suggesting a critical role of transcriptional regulation in the emergence of reversible therapy-resistance (i.e. persistence), a “bet-hedging” strategy to ensure survival in time-and/or space-varying environments (54).

Here, we have identified one such transcriptional programme in ER+ breast cancer cells that can cause a reversible resistance to the treatment of tamoxifen, an anti-estrogen therapeutic. Our results suggest that EMT and resistance to tamoxifen can drive each other; thus offering a mechanistic explanation for empirical observations showing that cells undergoing EMT are more resistant to tamoxifen (20), and that tamoxifen-resistant cells are EMT-like (21, 25). Notwithstanding this broad association between the two axes, which was validated in bulk and single-cell transcriptomic data, our model predicts the co-existence of and stochastic switching among six possible phenotypes – epithelial sensitive (ES), epithelial resistant (ER), hybrid sensitive (HS), hybrid resistant (HR), mesenchymal sensitive (MS), and mesenchymal resistant (MR) (**Fig 8A**). Thus, a progression to a full EMT need not be required for acquiring resistance to tamoxifen, and the association between at least a full EMT and drug resistance can be semi-independent, as seen for coupled EMT-stemness dynamics *in vitro* and *in vivo* (58, 59). Our stochastic simulations showed that cells can switch from an ES to MR state via HS or HR states, highlighting two alternative paths to acquire resistance. But whether cells show hysteresis in these paths (i.e. how reversible or irreversible can such transitions be) need to be investigated more carefully experimentally. Future efforts should focus on drawing more comprehensive landscapes for transitions along coupled axes of plasticity such as EMT, immune evasion, stemness and metabolic reprogramming (60), besides drug resistance. We also proposed SLUG as a marker for hybrid E/M and tamoxifen resistant phenotype, which may help explain association of worse clinicopathological features with high SLUG levels (35). While our model extensively features the connections among five players (ZEB1, SLUG, miR-200, ERα66 and ERα36) transcriptionally and their corresponding emergent phenotypes and dynamics, development of EMT and/or drug resistance is a much more complex process. Despite this limitation, the model simulations offer profound conceptual contributions. First, we postulate a putative mechanism that can explain diverse experimental observations and proposes a mutual dependence of two axes of plasticity that are crucial in cancer progression. Such models can help guide future experimental studies to investigate this coupling more comprehensively. Second, these specific molecular players can be thought of representative of various functional modules, and the fundamental feature of multistability demonstrated here can help conceptualize reversible phenotypic plasticity and non-genetic heterogeneity along multiple axes, as live-cell imaging and single-cell RNA-seq data becomes more prominent. Third, the population dynamics framework used here offers insights into how bidirectional plasticity and non-genetic heterogeneity can contribute to survival of a tumor cell population as well as the phenotypic demographic of the tumor colony.

**Figure 8:**
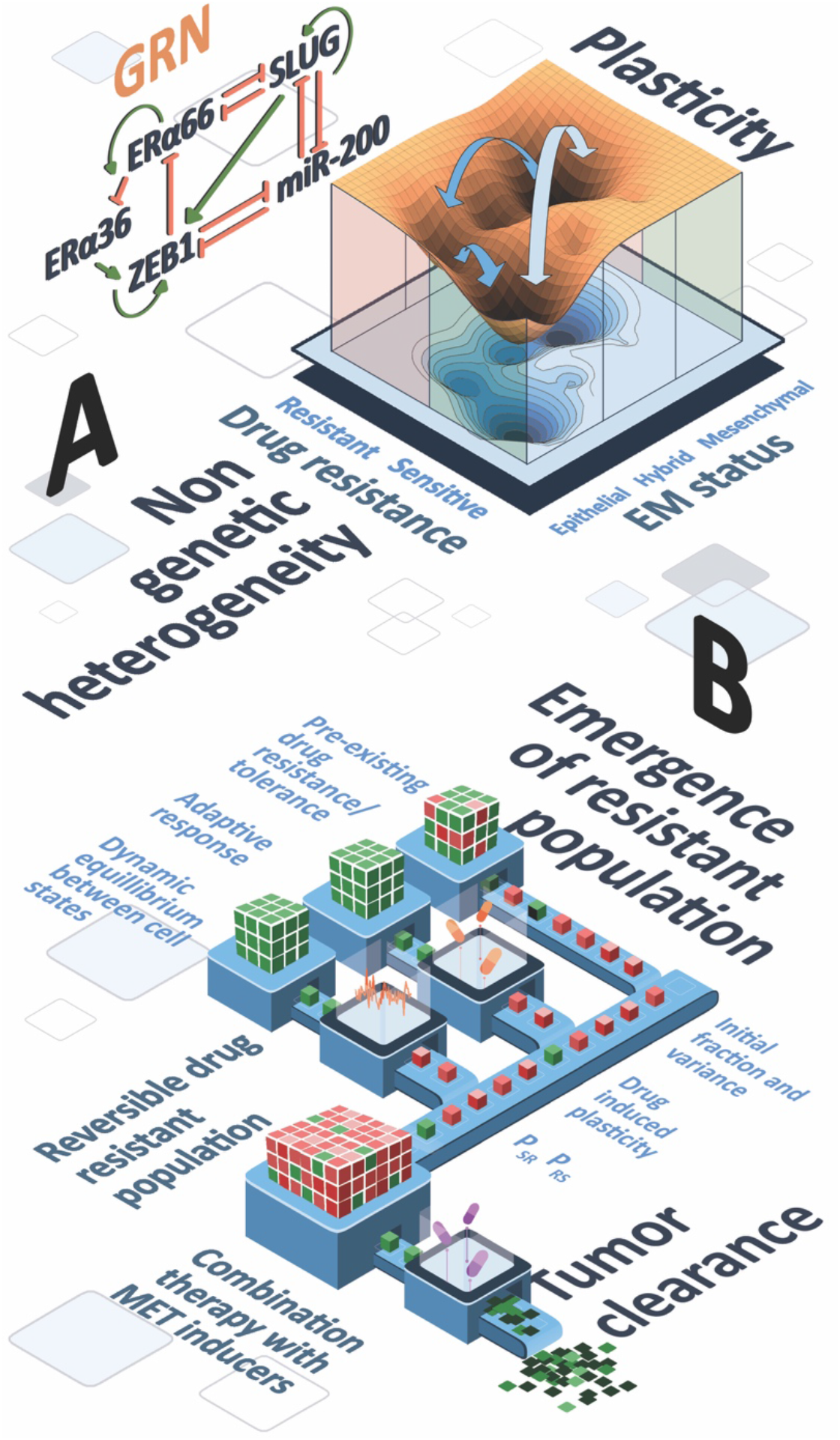
Schematic depicting dynamical traits of coupled EMT-ERα signaling network and its implications in tumor survival. **A**. Landscape showing multiple phenotypes defined on EMT and drug resistance axes: ES (epithelial-sensitive) and MR (mesenchymal-resistant) phenotypes are more dominant (witnessed by depth of the valley in the landscape). Arrows induced transitions under the influence of noise or drug among six phenotypes (ES, ER, HS, HR, MS, MR). **B**. Population dynamics showing multiple parallel paths to long-term resistance (pre-existing reversibly resistant cells, stochastic switching and dynamic equilibrium, and drug-induced plasticity) and the predicted effect of combinatorial therapy (anti-estrogen therapy + MET inducers) to drive population collapse.

The population dynamics framework has been maintained intentionally simple. Such simple frameworks are quite powerful in explaining many experimental phenomenon, as shown recently by Rehman *et al*. (51), where they could explain the maintenance of clonal complexity after drug treatment in tumors *in vivo* using a population dynamics model that assumes that all cells in the population are equally potent to switch to a reversibly resistant state and the choice of cell survival is completely independent of genetic background. One complexity that can be added to our model is the effect of ecological interactions between the two species – sensitive and resistant cells – such as cooperation or competition for resources (45–47). Other adaptation of the model can be to include cell-state transition probabilities as a function of relative stability of states calculated from the landscape estimated for an intracellular GRN. Similarly, death rate incorporated in our model can depend on internal challenge faced by cells during the metastatic cascade, such as anoikis and/or interaction with various immune cells.

Our population dynamics framework suggests that phenotypic plasticity (reversible transitions between drug-resistant and -sensitive phenotypes) and non-genetic heterogeneity (pre-existing reversibly resistant cells) can promote the survival of tumor population under drug treatment, with a higher contribution coming from plasticity, especially a switch from sensitive to resistant state. While heterogeneity aids in tumor survival, plasticity is crucial for the effects of heterogeneity to have a significant impact in terms of tumor fitness and survival. We found that the tumor size is maximum at an optimal plasticity level, beyond which the effect of plasticity, while still present, is reduced. These observations are consistent with reports about intermediate levels, not extremely high levels, of chromosomal instability being associated with the worst clinical outcomes (61). Furthermore, composition of final tumor population after a prolonged treatment was found to be variable from purely resistant to purely sensitive. Overall, the maintenance of long-term ‘resistance’ can be achieved via multiple paths: a) non-genetic heterogeneity in initial cell population, b) stochastic transitions among sensitive and resistant states driven by processes such as EMT, maintaining a dynamic equilibrium of cell subpopulations (30) and c) drug-induced transitions to a reversibly resistant state (**Fig 8B**).

Based on these features, we conceptually integrated the effect of MET on drug resistance, i.e. the broad association of an epithelial phenotype with a drug-sensitive state. We observed that MET inducers can minimize the impact of switch from sensitive to resistant phenotype, and can thus counteract the impact of both drug-induced plasticity and population heterogeneity to a large extent. Whether such combinatorial strategies are likely to be more effective simultaneously or sequentially (62, 63) is a question beyond the scope of this study, but can be answered by developing a multi-scale model including non-cell autonomous effects such as cooperation or competition, building on the principles of multistable regulatory networks and consequent phenotypic plasticity and non-genetic heterogeneity elucidated here. Future efforts to improve the predictability of such models will require rich dynamic experimental data based on which the observed cellular behavior can be realized in hyperspace of phenotypes obtained via mechanism-based and/or data-based investigation of regulatory networks.

## Acknowledgements

MKJ was supported by Infosys Foundation, Bangalore and by the Ramanujan Fellowship supported by Science and Engineering Research Board (SERB), Department of Science and Technology (DST), Government of India (SB/S2/RJN-049/2018). We would like to acknowledge Mr. Atchuta Srinivas Duddu for artwork (**Fig 3I, Fig 8**).

## Author contributions

MKJ designed and supervised research; SS, AM, HK, KH, SM and SM performed research and analysed data; SS, AM, HK, KH and MKJ wrote the paper.

## Code availability

All codes for the manuscript are available at: https://github.com/csbBSSE/TamRes

## Materials and methods

All details of model construction and data analysis are given in the Supplementary Information, together with Supplementary Figures.

## References

1. Salgia R, Kulkarni P (2018) The genetic/non-genetic duality of drug “resistance.” Trends in Cancer 4(2):110–118.

2. Bell CC, Gilan O (2020) Principles and mechanisms of non-genetic resistance in cancer. Br J Cancer. doi:10.1038/s41416-019-0648-6.

3. Marine J-C, Dawson S-J, Dawson MA (2020) Non-genetic mechanisms of therapeutic resistance in cancer. Nat Rev Cancer 20:743–756.

4. Bi M, et al. (2020) Enhancer reprogramming driven by high-order assemblies of transcription factors promotes phenotypic plasticity and breast cancer endocrine resistance. Nat Cell Biol 22(6):701–715.

5. Sharma S V., et al. (2010) A Chromatin-Mediated Reversible Drug-Tolerant State in Cancer Cell Subpopulations. Cell 141(1):69–80.

6. Howard GR, Johnson KE, Rodriguez Ayala A, Yankeelov TE, Brock A (2018) A multi-state model of chemoresistance to characterize phenotypic dynamics in breast cancer. Sci Rep 8:12058.

7. Pisco AO, et al. (2013) Non-Darwinian dynamics in therapy-induced cancer drug resistance. Nat Commun 4:2467.

8. Liau BB, et al. (2017) Adaptive Chromatin Remodeling Drives Glioblastoma Stem Cell Plasticity and Drug Tolerance. Cell Stem Cell 20(2):233-246.e7.

9. Su Y, et al. (2017) Single-cell analysis resolves the cell state transition and signaling dynamics associated with melanoma drug-induced resistance. Proc Natl Acad Sci U S A 114(52):13679–13684.

10. Musgrove EA, Sutherland RL (2009) Biological determinants of endocrine resistance in breast cancer. Nat Rev Cancer 9(9):631–643.

11. Burns KA, Korach KS (2012) Estrogen receptors and human disease: An update. Arch Toxicol 86(10):1491–1504.

12. Gonzalez-Angulo AM, Morales-Vasquez F, Hortobagyi GN (2007) Overview of resistance to systemic therapy in patients with breast cancer. Adv Exp Med Biol 608:1–22.

13. Shi L, et al. (2009) Expression of ER-α 36, a novel variant of estrogen receptor α, and resistance to tamoxifen treatment in breast cancer. J Clin Oncol 27(21):3423–3429.

14. Zhang XT, Wang ZY (2013) Estrogen receptor-α variant, ER-α36, is involved in tamoxifen resistance and estrogen hypersensitivity. Endocrinology 154(6):1990–8.

15. Yin L, Zhang X-T, Bian X-W, Guo Y-M, Wang Z-Y (2014) Disruption of the ER-α36-EGFR/HER2 Positive Regulatory Loops Restores Tamoxifen Sensitivity in Tamoxifen Resistance Breast Cancer Cells. PLoS One 9(9):e107369.

16. Wang ZY, et al. (2006) A variant of estrogen receptor-α, hER-α36: Transduction of estrogen-and antiestrogen-dependent membrane-initiated mitogenic signaling. Proc Natl Acad Sci U S A 103(24):9063–9068.

17. Kang L, et al. (2010) Involvement of Estrogen Receptor Variant ER-α36, Not GPR30, in Nongenomic Estrogen Signaling. Mol Endocrinol 24(4):709–721.

18. Wang Q, et al. (2018) Tamoxifen enhances stemness and promotes metastasis of ERα36 + breast cancer by upregulating ALDH1A1 in cancer cells. Cell Res 28:336– 358.

19. Zou Y, Ding L, Coleman M, Wang Z (2009) Estrogen receptor-alpha (ER-α) suppresses expression of its variant ER-α36. FEBS Lett 583(8):1368–1374.

20. Tian M, Schiemann WP (2017) TGF-β stimulation of EMT programs elicits non-genomic ER-α activity and anti-estrogen resistance in breast cancer cells. J Cancer Metastasis Treat 3(8):150.

21. Yuan J, et al. (2015) Acquisition of epithelial-mesenchymal transition phenotype in the tamoxifen-resistant breast cancer cell: A new role for G protein-coupled estrogen receptor in mediating tamoxifen resistance through cancer-associated fibroblast-derived fibronectin and β1-. Breast Cancer Res 17:69.

22. Zheng X, et al. (2015) Epithelial-to-mesenchymal transition is dispensable for metastasis but induces chemoresistance in pancreatic cancer. Nature 527(7579):525–530.

23. Wang Z, et al. (2009) Acquisition of epithelial-mesenchymal transition phenotype of gemcitabine-resistant pancreatic cancer cells is linked with activation of the notch signaling pathway. Cancer Res 69(6):2400–7.

24. Al Saleh S, Al Mulla F, Luqmani YA (2011) Estrogen Receptor Silencing Induces Epithelial to Mesenchymal Transition in Human Breast Cancer Cells. PLoS One 6(6):e20610.

25. Hiscox S, et al. (2006) Tamoxifen resistance in MCF7 cells promotes EMT-like behaviour and involves modulation of β-catenin phosphorylation. Int J Cancer 118(2):290–301.

26. Jiang Y, et al. (2014) Snail and Slug mediate tamoxifen resistance in breast cancer cells through activation of EGFR–ERK independent of epithelial–mesenchymal transition. J Mol Cell Biol 6(4):352–354.

27. Scherbakov AM, Andreeva OE, Shatskaya VA, Krasil’nikov MA (2012) The relationships between snail1 and estrogen receptor signaling in breast cancer cells. J Cell Biochem 113(6):2147–2155.

28. Zhang J, et al. (2017) ZEB1 induces ER-α promoter hypermethylation and confers antiestrogen resistance in breast cancer. Cell Death Dis 8(4):e2732.

29. Chamard-Jovenin C, et al. (2015) From ERaα66 to ERaα36: A generic method for validating a prognosis marker of breast tumor progression. BMC Syst Biol 9:28.

30. Jolly MK, Celia-Terrassa T (2019) Dynamics of Phenotypic Heterogeneity Associated with EMT and Stemness during Cancer Progression. J Clin Med 8(10):1542.

31. Castles CG, Oesterreich S, Hansen R, Fuqua SAW (1997) Auto-regulation of the estrogen receptor promoter. J Steroid Biochem Mol Biol 62(2–3):155–163.

32. Ye Y, et al. (2010) ERα signaling through slug regulates E-cadherin and EMT. Oncogene 29:1451–1462.

33. Li Y, et al. (2015) Slug contributes to cancer progression by direct regulation of ERα signaling pathway. Int J Oncol 46(4):1461–1472.

34. Thiebaut C, et al. (2017) Mammary epithelial cell phenotype disruption in vitro and in vivo through ERalpha36 overexpression. PLoS One 12(3):e0173931.

35. Subbalakshmi AR, Sahoo S, Biswas K, Jolly MK (2021) A computational systems biology approach identifies SLUG as a mediator of partial Epithelial-Mesenchymal Transition (EMT). Cells Tissues Organs:in press.

36. Huang B, et al. (2017) Interrogating the topological robustness of gene regulatory circuits by randomization. PLoS Comput Biol 13(3):e1005456.

37. Zhang X, et al. (2011) A Positive Feedback Loop of ER-α36/EGFR Promotes Malignant Growth of ER-negative Breast Cancer Cells. Oncogene 30(7):770–780.

38. Zhou C, et al. (2012) Proteomic analysis of acquired tamoxifen resistance in MCF-7 cells reveals expression signatures associated with enhanced migration. Breast Cancer Res 14:R45.

39. Chakraborty P, George JT, Tripathi S, Levine H, Jolly MK (2020) Comparative Study of Transcriptomics-Based Scoring Metrics for the Epithelial-Hybrid-Mesenchymal Spectrum. Front Bioeng Biotechnol 8:220.

40. Nakamura R, et al. (2018) Reciprocal expression of slug and snail in human oral cancer cells. PLoS One 13(7):e0199442.

41. Gras B, et al. (2014) Snail family members unequally trigger EMT and thereby differ in their ability to promote the neoplastic transformation of mammary epithelial cells. PLoS One 9(3). doi:10.1371/journal.pone.0092254.

42. Singh S, et al. (2021) Pan-cancer drivers are recurrent transcriptional regulatory heterogeneities in early-stage luminal breast cancer. Cancer Res:in press.

43. Cook DP, Vanderhyden BC (2020) Context specificity of the EMT transcriptional response. Nat Commun 11:2142.

44. Balázsi G, van Oudenaarden A, Collins JJ (2011) Cellular Decision Making and Biological Noise: From Microbes to Mammals. Cell 144(6):910–925.

45. Craig M, et al. (2019) Cooperative adaptation to therapy (CAT) confers resistance in heterogeneous non-small cell lung cancer. PLoS Comput Biol. doi:10.1371/journal.pcbi.1007278.

46. Bacevic K, et al. (2017) Spatial competition constrains resistance to targeted cancer therapy. Nat Commun 8:1995.

47. Nam A, et al. (2020) Phenotypic switching and adaptive strategies of cancer cells in response to stress: insights from live cell imaging and mathematical modeling. bioRxiv:028472.

48. Prieto-Vila M, et al. (2019) Single-cell analysis reveals a preexisting drug-resistant subpopulation in the luminal breast cancer subtype. Cancer Res 79(17):4412–4425.

49. Pisco AO, Huang S (2015) Non-genetic cancer cell plasticity and therapy-induced stemness in tumour relapse: ‘What does not kill me strengthens me .’ Br J Cancer 112:1725–1732.

50. Hangauer MJ, et al. (2017) Drug-tolerant persister cancer cells are vulnerable to GPX4 inhibition. Nature 551:247–250.

51. Rehman SK, et al. (2021) Colorectal Cancer Cells Enter a Diapause-like DTP State to Survive Chemotherapy. Cell 184(1):226-242.e21.

52. Seghers AC, Wilgenhof S, Lebbe C, Nyens B (2012) Successful rechallenge in two patients with BRAF-V600-mutant melanoma who experienced previous progression during treatment with a selective BRAF inhibitor. Melanoma Res 22(6):466–472.

53. Hata AN, et al. (2016) Tumor cells can follow distinct evolutionary paths to become resistant to epidermal growth factor receptor inhibition. Nat Med 22(3):262–269.

54. Jolly MK, Kulkarni P, Weninger K, Orban J, Levine H (2018) Phenotypic Plasticity, Bet-Hedging, and Androgen Independence in Prostate Cancer: Role of Non-Genetic Heterogeneity. Front Oncol 8:50.

55. Shaffer SM, et al. (2017) Rare cell variability and drug-induced reprogramming as a mode of cancer drug resistance. Nature 546:431–5.

56. Kawakami R, et al. (2020) ALDH1A3-mTOR axis as a therapeutic target for anticancer drug-tolerant persister cells in gastric cancer. Cancer Sci 111(3):962–973.

57. Hong SP, et al. (2019) Single-cell transcriptomics reveals multi-step adaptations to endocrine therapy. Nat Commun 10:3840.

58. Grosse-Wilde A, et al. (2015) Stemness of the hybrid epithelial/mesenchymal state in breast cancer and its association with poor survival. PLoS One 10(5):e0126522.

59. Pastushenko I, et al. (2018) Identification of the tumour transition states occurring during EMT. Nature 556(7702):463–468.

60. Kang X, Wang J, Li C (2019) Exposing the Underlying Relationship of Cancer Metastasis to Metabolism and Epithelial-Mesenchymal Transitions. iScience 21:754– 772.

61. Birkbak NJ, et al. (2011) Paradoxical relationship between chromosomal instability and survival outcome in cancer. Cancer Res 71(10):3447–3452.

62. Gunnarssson EB, D. S, Leder K, Foo J (2020) Understanding the role of phenotypic switching in cancer drug resistance. J Theor Biol 490:110162.

63. Goldman A, et al. (2015) Temporally sequenced anticancer drugs overcome adaptive resistance by targeting a vulnerable chemotherapy-induced phenotypic transition. Nat Commun 6:6139.

