## Supplementary Figures & Methods for "A mechanistic model captures the emergence and implications of non-genetic heterogeneity and reversible drug resistance in ER+ breast cancer cells"

Running title: EMT and therapy resistance in ER+ breast cancer cells

Sarthak Sahoo[1,2], Ashutosh Mishra[1,2,#], Harsimran Kaur[2,#], Kishore Hari[1], Srinath Muralidharan[3], Susmita Mandal[1], Mohit Kumar Jolly[1,*]

[1] Centre for BioSystems Science and Engineering, Indian Institute of Science, Bangalore, India
[2] Undergraduate Programme, Indian Institute of Science, Bangalore, India
[3] Department of Biotechnology, Indian Institute of Technology Madras, Chennai, India

### Supplementary Information

#### Materials and Methods

##### Random Circuit Perturbation Method (RACIPE)

**Simulations:**
Random Circuit Perturbation (RACIPE) (1) is a computational tool that generates an ensemble of kinetic models for a gene regulatory network given as input, simulated for a range of biologically relevant parameters and initial conditions. The input network is composed of inhibitory and activating links between each node. The expression of each node in the network is calculated through a set of Ordinary Differential Equations (ODEs) defined as follows:

$$\frac{dX_i}{dt} = g_{X_i} \prod_{j} H^s\left(X_j, X_{ji}0, n_{ji}, \lambda_{ji}\right) - k_{X_i} X_i$$

Here, $X_i, i \in \{1,2,3,4,5\}$ is the concentration of the nodes in the network, g is the basal production rate, k is the basal degradation rate and Hs is the shifted hill function that takes in the activator/inhibitory links into account to determine the production rate for the node. The parameters corresponding to each regulatory link are λ (fold-change parameter), n (Hill's coefficient) and X0 (threshold value for the Hill's function). The given network was simulated for 100000 parameter sets with the default settings for sampling of parameters.

**Normalization of the steady states generated through simulations:**
The generated output of steady states from RACIPE is in log2 form, hence to perform further analysis on the data, we performed z-score normalization, where expression values for each node are collated together. Mean and standard deviation is calculated for expression values of each node and then used to perform z-score normalization with the given formula.

$$Z_i = \frac{X_i - \underline{\underline{X_i}}}{\sigma_i}$$

**Boundary condition for the determination of phenotypes:**
The difference between z-score normalized expression values for ZEB1 and mir200 was used as the EM score and the difference between z-score normalized expression values for Era36 and Era66 was used as the Resistance score. These scores were used to discern a phenotype to each steady cell state generated through RACIPE. To determine the phenotypes, boundary conditions were defined for both EM and Resistance score. These boundary conditions were calculated using the mean of Gaussians in the score distributions with the given formula in equation x. The boundary conditions calculated for EM is -0.97 and 1.03, i.e. samples with scores > 1.03 are mesenchymal, those with < -0.97 are epithelial, and those between -0.97

and 1.03 are binned as hybrid E/M. Similarly, Resistance scores above -0.085 corresponds to resistant phenotype, and those below it correspond to sensitive phenotype.

$$EM_1 = mean(H) - \frac{[mean(H) - mean(E)]}{2}$$

$$EM_2 = mean(H) + \frac{[mean(M) - mean(H)]}{2}$$

$$R = mean(S) + \frac{[mean(R) - mean(S)]}{2}$$

**Uniform Manifold Approximation and Projection (UMAP):**
UMAP is a non-linear dimensionality reduction technique that preserves the global structure of the data while maintaining the local clustering. It was used to visualize the z-normalized simulated data in a 2D plane using the Python package *umap-learn*. Parameters n_neighbours and min_dist are crucial for determining the final structure of the data in the projection: n_neighbours controls the number of nearest neighbours required to construct initial high dimensional graph; min_dist focuses on the tightness of the Umap clusters. These parameters were set to 100 and 0.1 respectively to generate UMAP plots; colour scheme of the plot was decided by EM and Resistance score.

**Network Randomization:**
We created an ensemble of randomized "hypothetical" networks that are possible using the following rules:

(a) for each node in the "wild type" network, in each instance of randomization of the wild type network, the in-degree and the out-degree was kept fixed
(b) The number of activation and inhibitory edges in the entire network were kept fixed.
(c) Furthermore, the source node and the target node for each of the edges were kept constant, but the identity of the edge in terms of it being an activation or inhibition link was allowed to change.

RACIPE simulations were run on 100 such hypothetical networks. After downstream analysis as described above, a distribution of the correlation coefficients between the EM and the resistance score was computed.

**Network Perturbations:**
RACIPE generates an ensemble of models by randomizing the kinetic parameters within a range that also allows investigating the effect of network perturbations, such as over-expression and down-expression of a node. These perturbations were performed on the network by over/down expressing ZEB1 and Era66 by 50-fold using RACIPE. This implies that the range of production rate sampling is scaled by 50 times while down expression reduces the range of production rate sampling by 50 times. The percentage of each phenotype was calculated for each of the perturbation and plotted as a bar plot in comparison to the wild type to quantify its effect.

**K-means clustering and average Silhouette widths**

K-means clustering was employed on the z-normalized gene expression values with silhouette analysis to identify the optimal number of clusters in the data. Silhouette coefficient is a measure of the proximity between each point in a cluster to the each point in the neighboring clusters. The value near to 1 that the points in the cluster are far away from the points in the neighboring cluster whereas a value of 0 indicates an overlap between the clusters. *Scikit learn* packages K-means and Silhouette coefficient were used to estimate these values and a box plot was plotted corresponding to the number of clusters ranging from 2 to 8.

**Statistical testing**
We computed the Spearman correlation coefficients and used corresponding p-values to gauge the strength of correlations. For statistical comparison between groups, we used a two-tailed Student's t-test under the assumption of unequal variances and computed significance.

**Stochastic simulations**
We simulated the network using the Euler-Maruyama method for the parameters that showed the co-existence of 3 phenotypes: {ES, HR, MR} or {ES, HS, MR}. The corresponding equation is as follows:

$$X_i(t+1) = X_i(t) + \Delta t * g_{X_i} * \prod_j H^s\big(X_j(t), X_{ji}0, n_{ji}, \lambda_{ji}\big) - k_{X_i} * X_i(t) * \Delta t + \sqrt{\Delta t} * N(0,1)$$

The equation is just a discrete form of the ODE presented before, with an addition of the noise term $\sqrt{\Delta t} * N(0,1)$, where $\Delta t$ is the time step and $N(0,1)$ is a normal random variable with mean 0 and standard deviation 1. For each parameter set, we simulated the network for 100 different initial conditions sampled uniformly from the range $\left[0, 1.5 * \frac{g_{X_i}}{k_{X_i}}\right]$. We then normalized the trajectories using the mean and standard deviation of each node expression obtained from RACIPE and converted the trajectories to EM and Resistance scores in order to classify them into the observed phenotypes.

Using these trajectories, we constructed obtained a probability density (P) of the EM-Resistance score pairs and constructed a potential landscape by calculating the pseudo potential as $-log\ (P)$ (2). To demonstrate switching between the phenotypes, we simulated the network at the same parameter sets with the modified noise term: $n * X_i(t) * \sqrt{\Delta t} * N(0,1)$. Multiplication by $X_i(t)$ scales the noise to the levels to the node concentration and makes switching feasible.

**Gene expression data analysis (Clinical and single cell data)**

Publicly available microarray datasets were obtained from GEO and Spearman correlation coefficients were calculated for given genes. Single-sample gene set enrichment analysis (ssGSEA) (3) was performed on Hallmark EMT, Early Estrogen Response and Late Estrogen Response gene signatures obtained from MSigDB (Molecular Signatures Database) (4). TamRes signature was considered to be the set of upregulated genes obtained via proteomic analysis on resistant MCF7 cells (5). EMT scoring methods – 76GS, KS and MLR (6) – were used to compute the EMT scores for bulk microarray datasets. Higher 76GS scores indicate an epithelial phenotype, but higher KS and MLR scores denote a mesenchymal phenotype. Thus, usually, 76GS scores correlate negatively with KS and MLR scores.

Activity values computation for 10-cell and single cell datasets were done using AUCell (7). The BRCA ESR1 specific regulon was obtained from the database GRNDb (8). Signatures for epithelial and mesenchymal phenotypes (cell lines) were taken from Tan *et al.* (9).

**Population dynamics**

**Model Formulation:**
The population dynamics model that we constructed consists of cells that have been modelled explicitly with each cell having its attributes during the simulation. The three main processes during the growth/decline of a population of cells that we modelled are:

(a) proliferation,
(b) death, and
(c) switching between states.

We consider that the cells would be present in one of two states – Sensitive (S) or Resistant (R). We defined a parameter called the "resistance score". The resistance score is a number that represents the fitness of a cell in the presence of an anti-estrogen drug. The score varies in an arbitrary range of -6 to +6. Cells with smaller values of "resistance score" are likely to be more sensitive to the drug than cells with higher scores. We incorporated variability into the system (representative of heterogeneity) by sampling the resistance scores which are assigned to each cell from a gaussian distribution centered a fixed value of mean but with different standard deviations. Each cell is assigned with 2 scores, one each for its possible sensitive (S) or resistant (R) state, former sampled from a gaussian centered at -2 and with a fixed variance and the latter sampled from a gaussian centered at +2 and with a fixed variance. These scores are assigned during the birth of the cell and remain fixed over the course of the simulation. Each cell has an index variable that keeps track of the current status of the cell – S or R. Depending on the current status of the cell, the probability of the death of cell due to the drug varies depending on the corresponding "sensitive" or "resistant" score.

Cell death can occur through 2 independent ways – a constant basal probability of cell death and a death due to the presence of the drug. The constant basal probability of cell death is a fixed number kept constant (at 0.1) over all simulations unless specified. The probability for drug induced cell death is dependent on the current status of the cell and the corresponding score assigned to it during its birth. To map the resistance score of each cell to a probability of death, we used a sigmoidal function. So, if a cell has a resistance score of x, then the probability of its death is given by $\frac{e^x}{e^x + c}$, where c is a constant. For our simulations, we used c = 0.6. Varying c doesn't change the qualitative observations of our model.

At a population level, we assume a logistic growth model for the cell population. Cells are allowed to proliferate with a probability given by:

$$Proliferation\ rate * \left(1 - \frac{current\ population\ size}{carrying\ capacity}\right)$$

Proliferation rate was set at 0.91 unless specified, and carrying capacity was set at 10^5 cells. For our model, we assumed that the proliferation and the basal death rates of the cells were same irrespective of the current state of the cell. Upon cell division, the daughter cells retain the status of the mother cell. So, if a cell was sensitive at the time of cell division, then both the daughter cells would also be initialized to be sensitive. However, to account for variability during cell division, the 2 scores assigned to each cell are resampled from the gaussian distributions with the specified variance (heterogeneity level).

Cells can also switch between sensitive and resistant states. Transition probability from sensitive to resistant state is given by $P_{SR}$; that from resistant to sensitive state is given by $P_{RS}$. These probabilities characterize the plasticity of the system. The basal death probability, probability of proliferation, heterogeneity, and transition probability constants are kept the same for a single simulation. Later, we perform a set of simulations by changing each of these parameters individually to show their effect at the population level.

**Simulation:**

For the simulation, we start with an initial population of 100 sensitive cells, with scores distributed according to the level of heterogeneity in the system (specified as a given value of std. dev. of gaussians from which the scores are sampled from). Then we track the dynamics of this population over the course of 100 timesteps. Each cell gets the chance to go through all three events (proliferation, death, switching) once in a timestep. Given the typical timescale of cell division in many mammalian systems (~24-30 hours), and the proliferation probability considered here (=0.91), each timestep can be conceptually mapped on to one day.

To reduce the bias of choosing the cells in a particular order, we shuffle the order of sequence of the above events. If the cell proliferates, then the daughter cells would be allowed to do all these three actions on the next day. Furthermore, if the cell dies before proliferating/switching, then it would not be allowed to proliferate or switch further. Additionally, we monitor the sum of the current number of cells and the progenies that they have produced, and we don't allow cells to proliferate if this number has reached the carrying capacity. This condition is important to ensure that the total number of cells does not exceed the carrying capacity at any point in time.

Finally, we create a summary of this simulation in which we find out the diversity of the population by distributing the population into bins of size 0.1 according to their resistance score. As we are not considering cooperation and/or competition in our model, sensitive cells are more likely to die in the presence of the drug while the dynamics of resistant cells are less affected by the presence or absence of the drug.

**Initial fraction of resistant cells:**
In this case, we vary the initial condition. Instead of starting with all sensitive cells, we start with a fraction of initial population as resistant cells. This way we can start with heterogenous population and we find out the effects on the dynamics of the population. For initial fraction x, x out of 100 cells are sampled from the Gaussian corresponding to resistant cells (resistance score > 0). Additionally, we define an "extinction probability" which measures the fraction of cases in which a cell population collapses for a particular value of plasticity and heterogeneity. This statistics can help to find the "critical cases" and characterize how parameters like *intrinsic* heterogeneity or heterogeneity in initial population affects the dynamics.

**Drug induced plasticity & MET induced sensitivity:**
In this case, we study how plasticity due to the presence of drug affects the overall dynamics of the population. Here, we introduce a parameter called "Drug induced plasticity" (DIP), a probability value based on which susceptible cells can switch to resistant cells. Similarly, we have another probability value for the presence of MET ("MET-induced sensitivity" - MIS), under which resistant cells can switch to sensitive cells. We can vary the parameters DPI and MIS individually or simultaneously and see how these affect the final population size and population distribution.

DIP and MIS are defined independently of P_SR and P_RS. Thus, the effective probability of switching from sensitive to resistant would be
P_SR + (1 – P_SR)*DIP

Similarly, For MET, effective switching probability from resistant to sensitive would be

P_RS + (1 – P_RS) * MIS

**References**


1. Huang B, et al. (2017) Interrogating the Topological Robustness of Gene Regulatory Circuits. *PLoS Comput Biol* 13(3):e1005456.

2. Wang J, Zhang K, Xu L, Wang E (2011) Quantifying the Waddington landscape and biological paths for development and differentiation. *Proc Natl Acad Sci U S A* 108:8257–8262.

3. Subramanian A, et al. (2005) Gene set enrichment analysis: A knowledge-based approach for interpreting genome-wide expression profiles. *Proc Natl Acad Sci U S A* 102(43):15545–15550.

4. Liberzon A, et al. (2011) Molecular signatures database (MSigDB) 3.0. *Bioinformatics* 27(12):1739–1740.

5. Zhou C, et al. (2012) Proteomic analysis of acquired tamoxifen resistance in MCF-7 cells reveals expression signatures associated with enhanced migration. *Breast Cancer Res* 14:R45.

6. Chakraborty P, George JT, Tripathi S, Levine H, Jolly MK (2020) Comparative study of transcriptomics-based scoring metrics for the epithelial-hybrid-mesenchymal spectrum. *Front Bioeng Biotechnol* 8:220.

7. Aibar S, et al. (2017) SCENIC: Single-cell regulatory network inference and clustering. *Nat Methods* 14:1083–1086.

8. Fang L, et al. (2021) GRNdb: Decoding the gene regulatory networks in diverse human and mouse conditions. *Nucleic Acids Res* 49(D1):D97–D103.

9. Tan TZ, et al. (2014) Epithelial-mesenchymal transition spectrum quantification and its efficacy in deciphering survival and drug responses of cancer patients. *EMBO Mol Med* 6(10):1279–1293.


#### Supplementary Figures

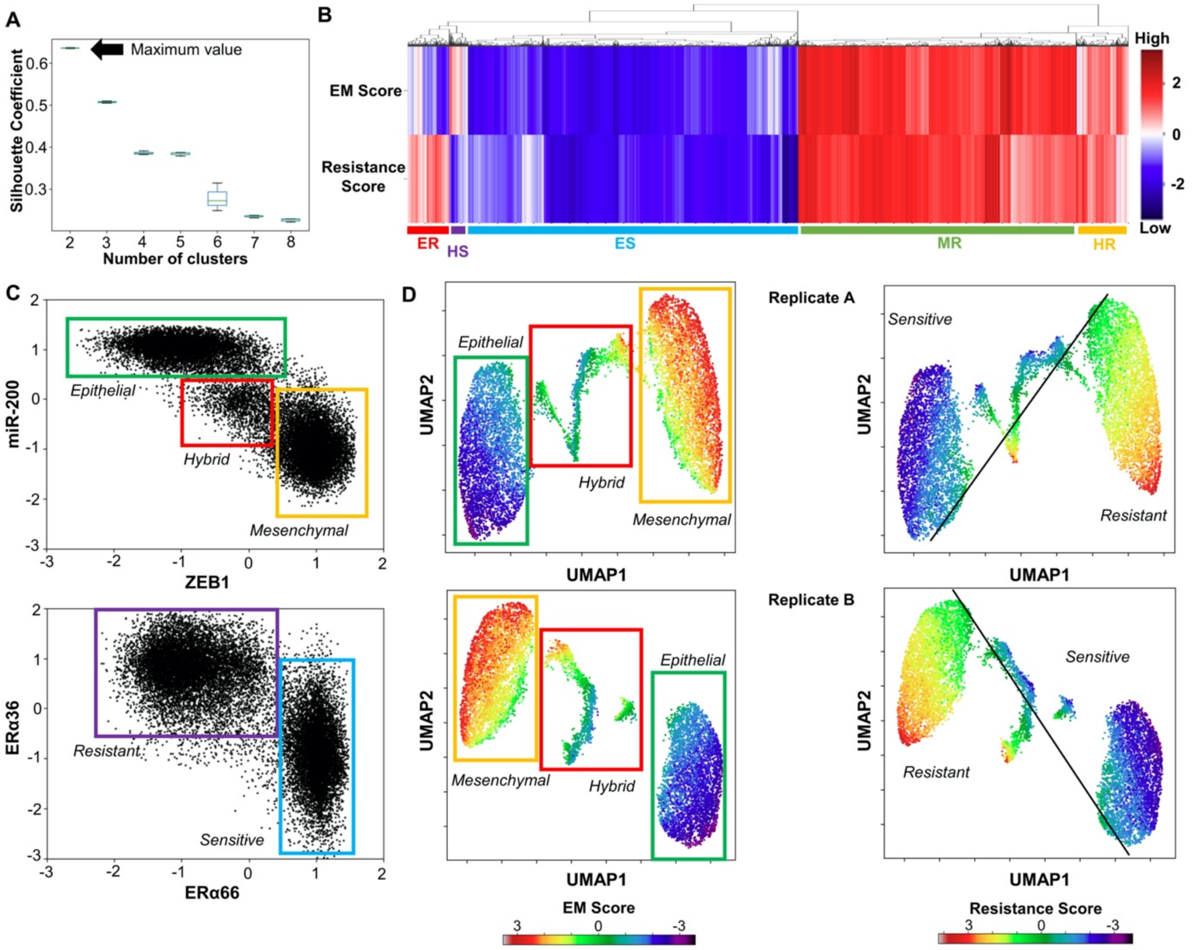


***Supplementary Figure S1: Characterization of EM and drug resistant phenotypes. A.*** *Average silhouette coefficient values for different number of clusters (2 to 8). Maximum value at number of clusters = 2 indicates the presence of 2 primary clusters in the data.* ***B.*** *Heatmap of EM scores and Resistance scores with cluster dendrogram revealing the existence multiple phenotypes (HS, ER and HR) other than the primary ES and MR phenotypes.* ***C.*** *Scatter plot*

*of all RACIPE solutions on ZEB1-miR200 plane, and on ERα66 and ERα36 plane.* ***D.*** *Two additional replicates showing the reproducibility of UMAP dimensionality reduction plots on all steady states obtained by RACIPE colored by either the EM Score or the Resistance Score.*

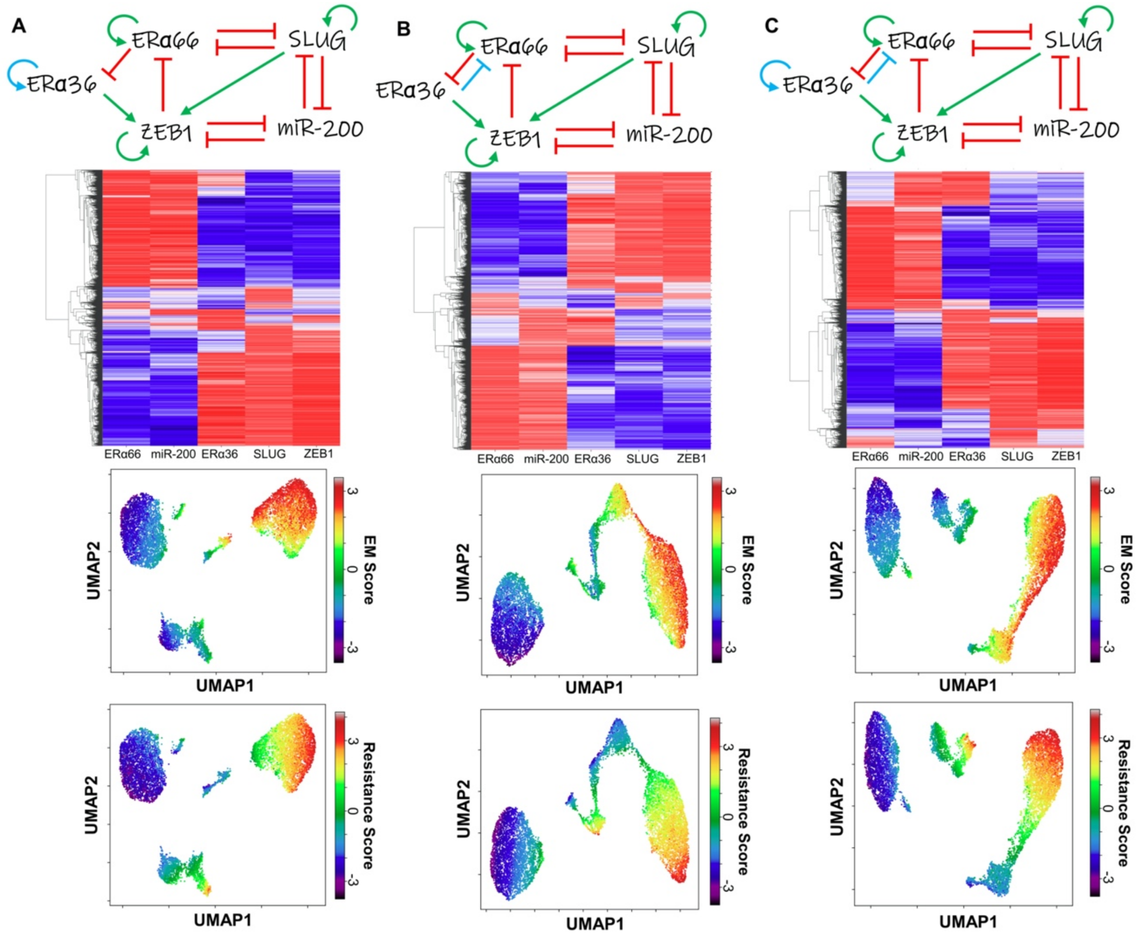


***Supplementary Figure S2: Alternative GRN simulations considering additional links. A.*** *Extra link of self-activation considered for ERα36 (shown in blue).* ***B.*** *Extra link for the inhibition of ERα66 by ERα36 (shown in blue)* ***C.*** *Both links considered. Clustered heatmaps with UMAP plots (colored by EM & resistance scores) are shown for each of the 3 corresponding cases.*

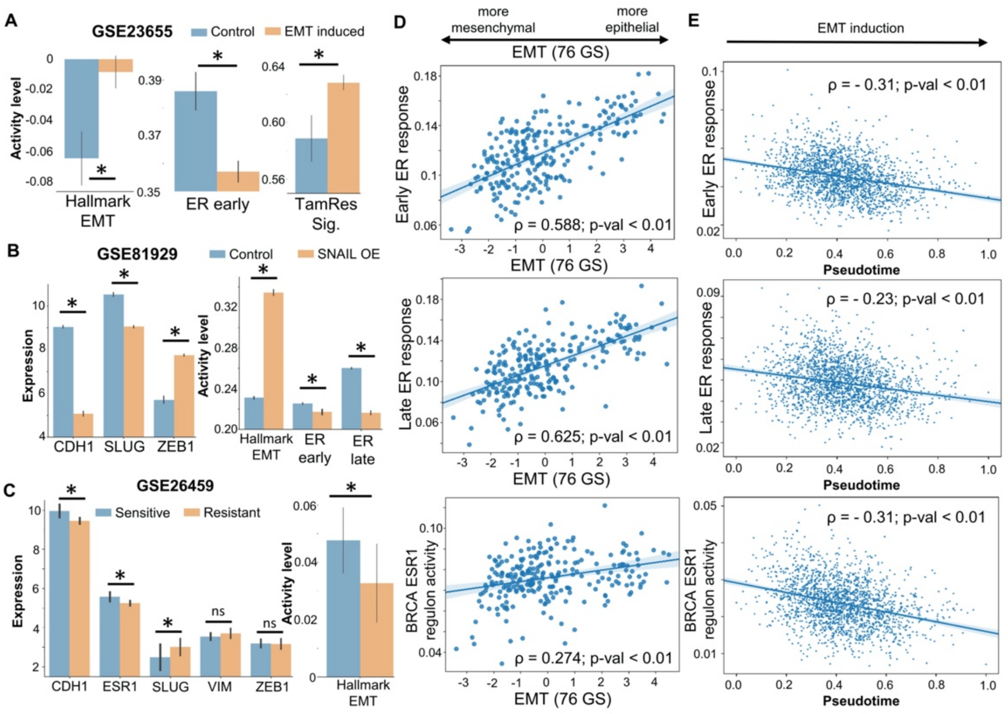


***Supplementary Figure S3: Additional analysis of publicly available experimental data.*** ***A.*** *Experimental data showing EMT induction via Six1 over expression in ER+ breast cancer MCF7 cells and concurrent decrease in magnitude of early estrogen response and an increase in Tamoxifen resistance signature (GSE23655).* ***B.*** *Experimental data showing EMT induction via Snail over expression MCF10A cells and the concurrent decrease in the magnitude of early and late estrogen response. Changes in gene expression levels of CDH1, SLUG and ZEB1 are shown too (GSE81929).* ***C.*** *Experimental data showing differences in expression levels of CDH1, VIM, ESR1, ZEB1 and SLUG and activity levels of EMT programme in sensitive and resistant MCF7 cell lines (GSE26459).* ***D.*** *Scatter plots showing associations with 76GS EMT scoring metric and the early, late estrogen response gene activity and BRCA ESR1 regulon activity (GSE147356).* ***E.*** *Scatter plots showing the change in early, late estrogen response gene activity and BRCA ESR1 regulon activity with pseudotime in the dataset GSE147405.*

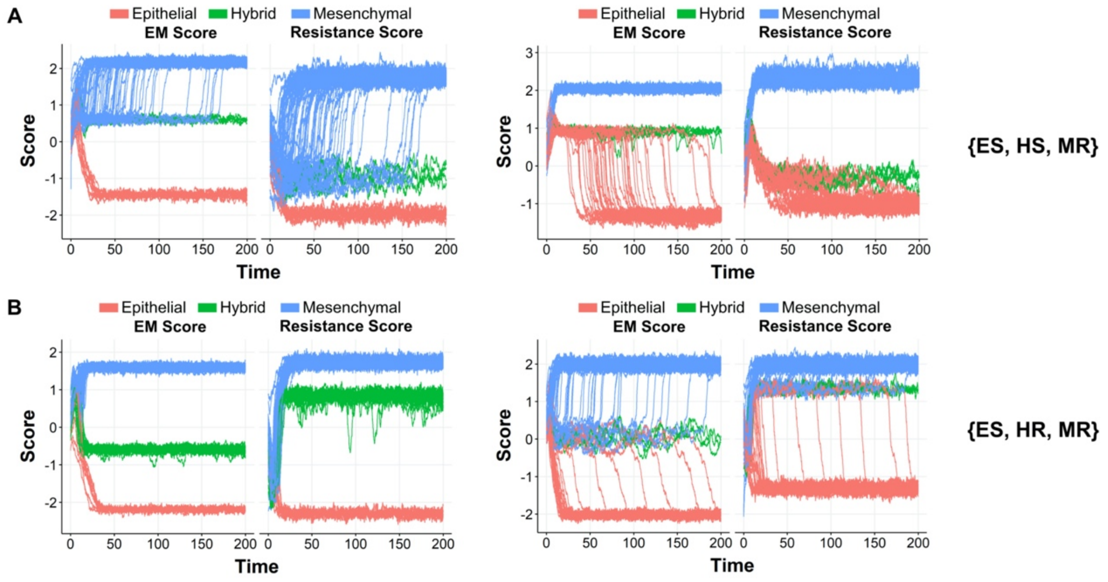


***Supplementary Figure S4: Stochastic simulations*** *Representative parameter sets from multiple initial conditions showing dynamic profiles in EM and resistance scores for the phases :* ***A.*** *{ES, HS, MR},* ***B.*** *{ES, HR, MR}*

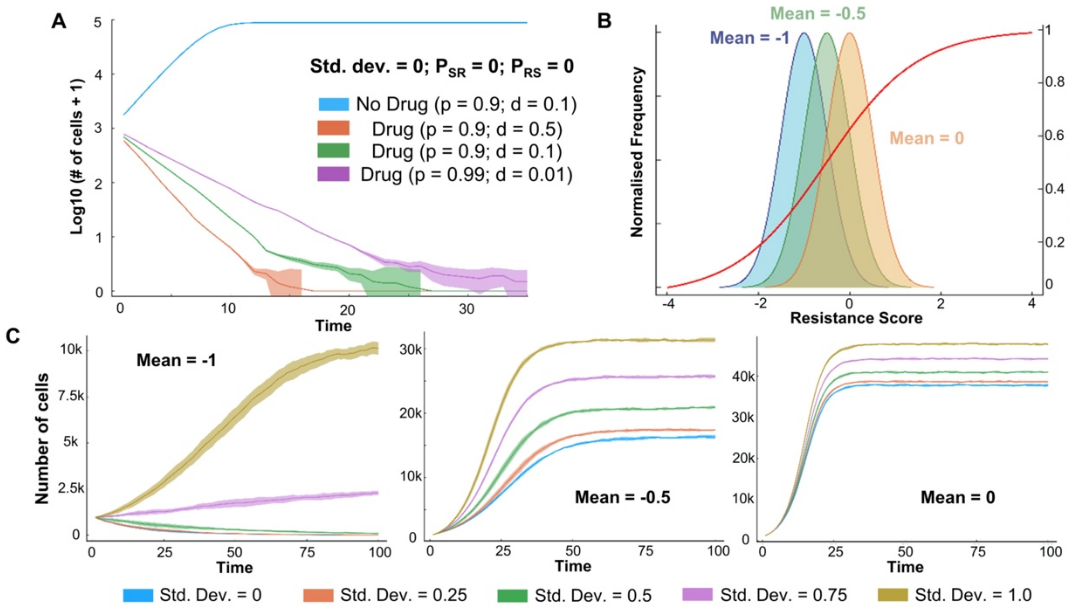


***Supplementary Figure S5: Population dynamics framework. A.*** *Effect of different proliferation and death probabilities on population sizes over time, when starting with an initially all sensitive cell population without any heterogeneity (Std. dev. = 0) or plasticity ($P_{SR}$ = $P_{RS}$ = 0).* ***B.*** *Schematic showing the distribution of a unimodal sampling strategy for a population having a single phenotype.* ***C.*** *Population sizes over time as a function of the different sampling strategies for Gaussian distributions (Mean at -1, -0.5 and 0) with different heterogeneity (std. dev. = 0, 0.25, 0.5, 0.75 and 1.0).*

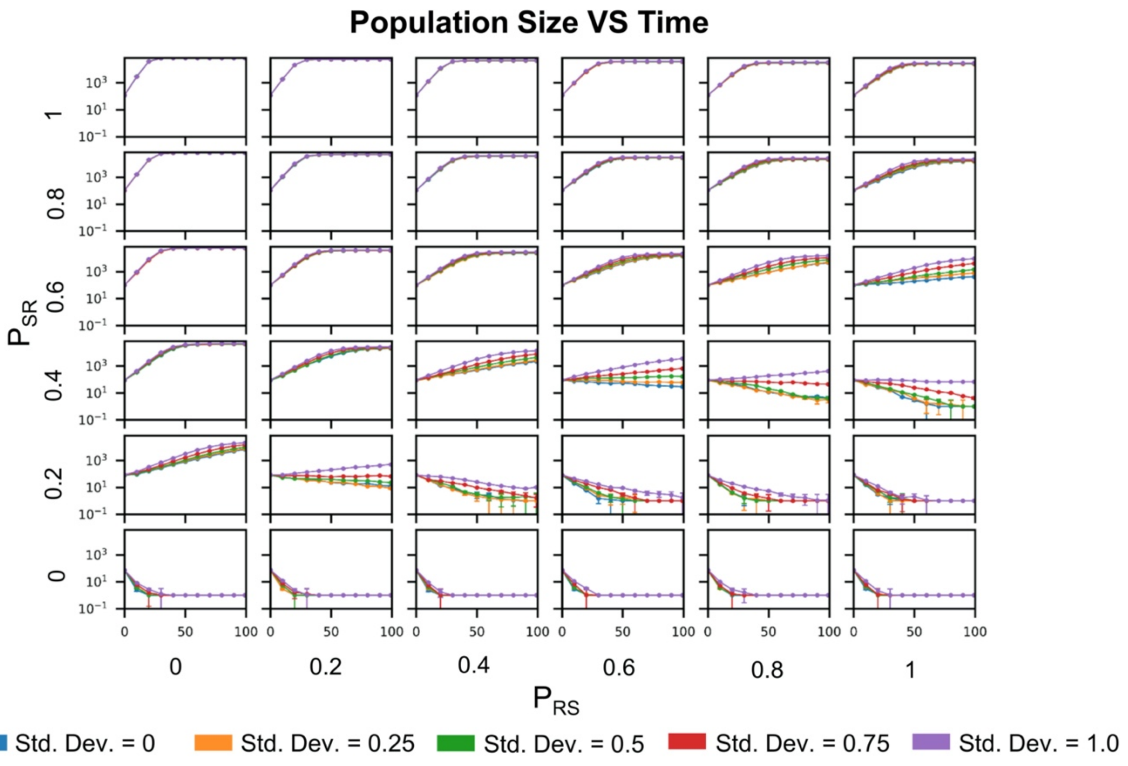


***Supplementary Figure S6: Population dynamics profiles.*** *Cell population sizes as a function of time, across multiple values of $P_{SR}$ and $P_{RS}$ with varying levels of heterogeneity (std. dev. = 0, 0.25, 0.5, 0.75 and 1.0)*

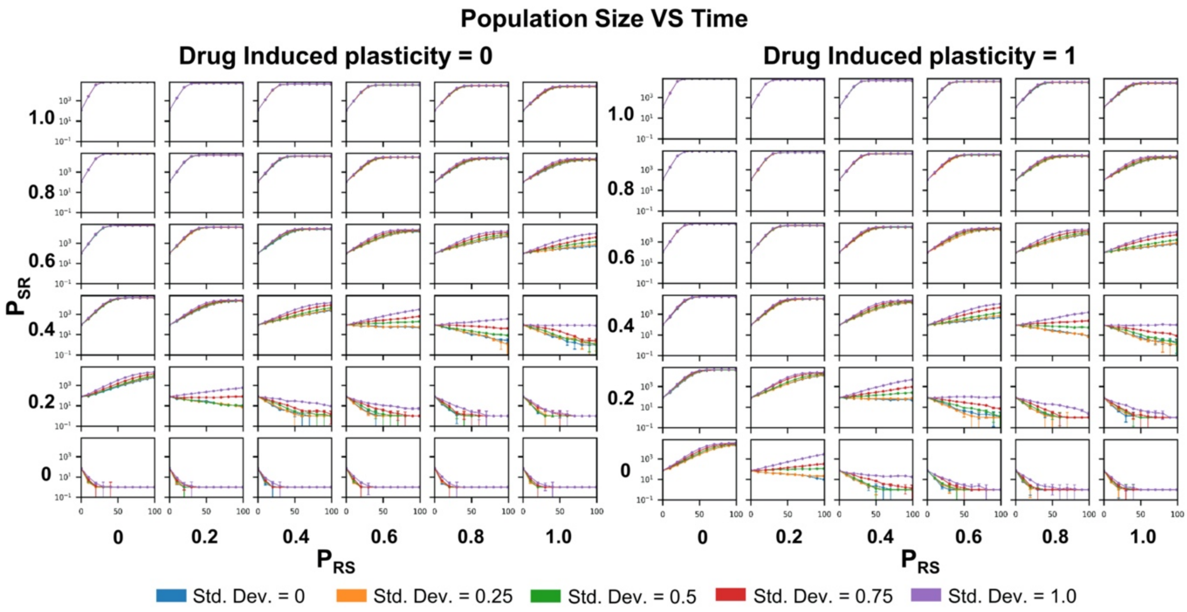


***Supplementary Figure S7: Population dynamics profiles with varied drug-induced plasticity.*** *Population sizes as a function of time across multiple values of $P_{SR}$ and $P_{RS}$ with varying levels of heterogeneity (std. dev. = 0, 0.25, 0.5, 0.75 and 1.0) at two extreme levels of drug induced plasticity (0 and 1).*
